# Nano-clusters of ligand-activated integrins organize immobile, signalling active, nano-clusters of phosphorylated FAK required for mechanosignaling in focal adhesions

**DOI:** 10.1101/2024.02.25.581925

**Authors:** Kashish Jain, Rida F. Minhaj, Pakorn Kanchanawong, Michael P. Sheetz, Rishita Changede

## Abstract

Transmembrane signalling receptors, such as integrins, organise as nanoclusters that are thought to provide several advantages including, increasing avidity, sensitivity (increasing the signal-to-noise ratio) and robustness (signalling above a threshold rather than activation by a single receptor) of the signal compared to signalling by single receptors. Compared to large micron-sized clusters, nanoclusters offer the advantage of rapid turnover for the disassembly of the signal. However, if nanoclusters function as signalling hubs remains poorly understood. Here, we employ fluorescence nanoscopy combined with photoactivation and photobleaching at sub-diffraction limited resolution of ∼100nm length scale within a focal adhesion to examine the dynamics of diverse focal adhesion proteins. We show that (i) subregions of focal adhesions are enriched in immobile population of integrin β3 organised as nanoclusters, which (ii) in turn serve to organise nanoclusters of associated key adhesome proteins-vinculin, focal adhesion kinase (FAK) and paxillin, demonstrating that signalling proceeds by formation of nanoclusters rather than through individual proteins. (iii) Distinct focal adhesion protein nanoclusters exhibit distinct dynamics dependent on function. (iv) long-lived nanoclusters function as signalling hubs-wherein phosphorylated FAK and paxillin formed stable nanoclusters in close proximity to immobile integrin nanoclusters which are disassembled in response to inactivation signal by phosphatase PTPN12 (v) signalling takes place in response to an external signal such as force or geometric arrangement of the nanoclusters and when the signal is removed, these nanoclusters disassemble. Taken together, these results demonstrate that signalling downstream of transmembrane receptors is organised as hubs of signalling proteins (FAK, paxillin, vinculin) seeded by nanoclusters of the transmembrane receptor (integrin).

## Introduction

Recent observations by various super-resolution microscopy modalities have shown that a wide range of transmembrane receptors, including integrins [1], cadherins [2], T-cell receptors [3, 4], immunological synapses [5], neuronal synapses [6], GPI-anchored proteins [7], organize into sub-micron sized clusters within larger signalling assemblies on the plasma membrane. These nanocluster assemblies can provide multiple advantages to bring about effective signalling including increasing local concentration of signalling, increasing signal-noise ratio compared to single molecule signalling, allowing rapid disassembly compared to large micron sized structures, and signal integration of a discrete signal (assuming each nanocluster can act as a signalling hub) to bring about the required final output [8–11]. However, this is a hypothesis at present, and scientific evidence to prove or disprove it is lacking [12]. One key signalling event that is the most well studied hallmark of signalling in biology, in response to external signals, is phosphorylation of kinases, e.g. force dependent phosphorylation of focal adhesion kinase (FAK) at focal adhesions on rigid substrates [13–16], or DAPK1 at podosomes on soft substrates [17], which in turn phosphorylate downstream signalling molecules and activates them. This signalling coordinates downstream cellular functions including cytoskeletal reorganisation, gene expression (regulated by translocation of activated LIM domain protein); paxillin translocation to the nucleus (activated by phosphorylation at the focal adhesions) [18, 19], or DAPK1-induced *anoikis* on soft substrates [17], among others. The dynamics of adhesions is closely regulated by force dependent activation of kinases such as FAK, and deactivation of signal by force dependent phosphatase-PTPN12, (also known as PTP-Pest) [13–16, 20–23]. The absence of PTPN12 function leads to the formation of large adhesions on compliant substrates lacking force [24]. The spatial organisation and regulation of these signal proteins required for signalling remains elusive. Do they function as single molecules or as a consort of nanoclusters?

Robust signalling depends on a fine balance between signal activation and deactivation in a precise spatio-temporal manner. Disruptions of this balance are hallmarks of many diseases including cancer and developmental deformities [25–28]. Signalling has to occur in a small but finite space (larger than a single receptor) for specific durations. Let us analyse the extreme cases: if signalling were solely dependent on individual receptor activation, then the signal-to-noise ratio would be poor, leading to many false positives. Conversely, if the entire micron-sized focal adhesion consisted solely of activated integrins, their rapid disassembly would be challenging, especially in situations requiring rapid signal cycling, such as during cell migration. Thus, similar to a gecko’s feet, where each individual foot pad provides strong adhesion force but can be detached quickly due to its small surface area, nanoclustering can provide various advantages over single receptor signalling, or large micron-scaled clusters [8]. Nanoclustering can provide a higher spatio-temporal stability compared to the freely diffusing single molecules and increase local concentration of the signalling molecules where they are protected from small deactivation events (e.g. by a single accidentally activated phosphatase) further increasing the robustness of the signal [8, 12]. In the case of integrin nanoclusters, formation of a nanocluster requires the presence of multiple activated integrins [29, 30]. Further, nanoclusters can regulate the time taken by these stop signals to act in concert with cellular function [24]. Moreover, these nanoclusters can act as discreet signalling events, which can be integrated to determine the overall strength of the signalling response by the whole cell. Without this spatial organisation, robust signalling would be challenging.

In this study, we aim to investigate whether there is functional organisation within focal adhesions and whether nanoclusters of integrins can serve as signalling hubs within these adhesions. If nanoclusters indeed function as signalling hubs, integrin nanoclusters would (i) be organised as nanoclusters (shown before (ii) which would persist in time compared to the neighbouring single molecules. Currently, single molecule studies have revealed that individual integrins can be immobile or mobile and the immobilisation of integrins is regulated by ECM and talin binding, and traction force [31, 32], however whether this immobile population is organised within nanoclusters is not known. Longer length scale (multiple micron size) population dynamics of focal adhesions have revealed that adhesion dynamics and maturation depend on cytoskeletal adhesion proteins such as FAK and vinculin, and on actin organisation [33–36]. (iii) they would organise downstream signalling molecules in nanoclusters which would also depict long lived dynamics (iv) disassemble rapidly when a signal is turned off and (v) not form immobile clusters when appropriate external signal is not present.

To test this hypothesis, instead of single molecules or large micron sized adhesions, we measured the immobilisation and life-time of diffraction-limited nanoclusters of integrins and key consensus adhesome proteins with diverse functions, using Fluorescence Loss After Photoactivation (FLAP) [22] and Fluorescence Recovery after Photobleaching (FRAP) [37] at nanoscale of multi-molecular assemblies and termed it nanoKymo-FLAP and nanoKymo-FRAP. Key adhesome proteins [38] include transmembrane receptors-integrins, that bind to the extracellular matrix and sense the force, key force dependent kinase-FAK, that can in turn phosphorylate and activate other downstream signalling proteins such as paxillin [13–16], force transducer-vinculin, and LIM domain protein-paxillin, that transmits the signal to the nucleus to regulate gene expression in response to force [38–40]. We tested if these proteins are organised as immobile nanoclusters within the adhesions, the relative positions of these nanoclusters with respect to the integrin nanoclusters. To understand the role of these nanoclusters and its dynamics in response to signalling, we studied it in response to change in signal using (i) mutants or (ii) downregulation of signal by eliminating force by ROCK inhibitor Y-27632, or (iii) changing the spatial organisation of nanoclusters using nanopatterning which does not allow robust adhesions to form [41]. We found that integrin nanoclusters organise long-lived activated FAK nanoclusters which bring about functional signalling directly in response to external cues and absence of these cues eliminates signalling by disassembly of the signalling hubs of nanoclusters.

## Results

### NanoKymo-FLAP resolves 2 distinct spatiotemporal populations of Integrin β3 nanoclusters

We investigated the organization and spatio-temporal dynamics of integrin β3 in focal adhesions. Mouse embryonic fibroblasts expressing β3-GFP, were spread on fibronectin-coated glass substrate for 15 minutes. After 15 minutes of cell spreading, we imaged live cells using super-resolution confocal microscopy, Live SR. Live SR (provided by Roper Scientific France) provides a ∼2-fold higher resolution over the conventional pinhole confocal microscopy by optically demodulated structured illumination combined with a spinning disc confocal microscopy [30, 42–45]. This is optimal to image structures close to the cover slip such as focal adhesions, as the increase in resolution is not as pronounced further away from the cover slip.

Imaging of integrins in early focal adhesions revealed regions of high and low intensity (Fig. 1a, Movie S1.). We characterised the integrin β3 nanocluster distribution by thresholding the integrin β3-GFP fluorescence images and performed nearest neighbour distance analysis of segmented integrin β3 clusters. An average centre-to-centre distance of 567.21±3.87 nm (mean±S.E.M.; Fig. 1b) was observed between these clusters, indicating that in focal adhesions integrin β3 are arranged with a characteristic distance between them [1, 41].

**Fig. 1.**
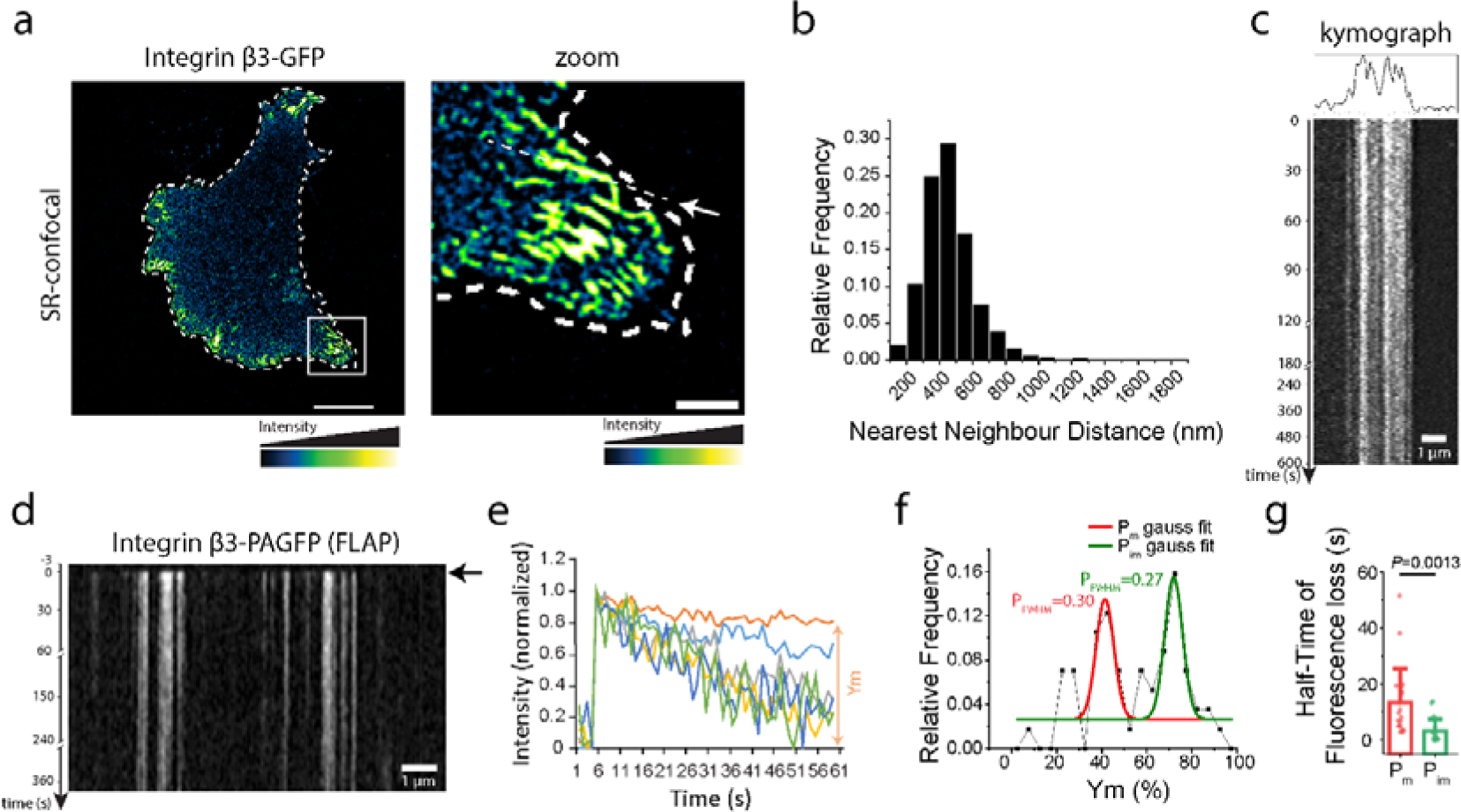
Integrin β3 organizes in nanoclusters with 2 distinct populations in focal adhesions. (a) Representative super-resolution confocal image of MEFs overexpressing integrin β3-GFP, along with cell boundary in white dashed line and zoom of the white box (right). Scale bar (left) 10 µm. Scale bar (right) 2µm. (b) Relative frequency of the Nearest neighbour distance of integrin β3 nanoclusters. N=2626 from 8 cells. (c) Kymograph of the dotted line in (a) along with line profile of the first frame on top. (d) Kymograph obtained from nanoKymo-FLAP of integrin β3-PAGFP with the black arrow showing photoactivation time. (e) Line profiles obtained from kymographs of nanoKymo-FLAP, showing normalized intensity vs time plot, along with an example of Ym. f) Relative frequency (histogram) of Ym calculated from intensity vs time plots, along with Gaussian fits showing 2 populations. P_FWHM_ denotes the fraction of population lying in the full width half maxima of the Gaussian fit. N=57 from 6 cells. (g) Mean±S.D. plot of half-time of fluorescence loss of the 2 populations, along with the individual points. P value was evaluated by two-tailed t-test.

Single molecule studies have revealed that individual integrins can be immobile or mobile, however population dynamics within nano-regions of focal adhesions has not been investigated. There are 3 possible dynamics of integrin nanoclusters, (i) they are random patches of high and low intensity which appear in a single snapshot and are not persistent over time, (ii) they are well defined nanoclusters which persist over time, where they form and disassemble as a cluster, or (iii) they are loosely defined nanoclusters which constantly exchange molecules with the surrounding this is indicated by Fujiwara et al., 2023 [46], using ultra high speed imaging of single molecules. However, using single molecule analysis lacks the resolution to analyse the combined population dynamics (nanocluster size ranging 13-100 nm), which we aim to address here. To understand behaviour of the nanoclusters, we investigated which dynamics appear over time and subsequently how the dynamics might relate to function.

To understand the dynamics of integrin β3 nanoclusters, we performed time lapse imaging over 10 minutes. Imaging was performed at intervals of 2 seconds initially (for 2 minutes) followed by 10 seconds (for 8 minutes, to prevent excess bleaching) and kymographs were plotted, in this case, along the length of focal adhesions (dotted line in figure 1a). We further analysed the kymographs to reveal the half-life and mobile versus immobile fraction i.e. Ym of the analysed protein. This Kymograph analysis of focal adhesions revealed that regions of high intensity persisted over time, indicating that integrin β3 in these nanoclustered regions were relatively immobile (Fig. 1b, Movie S1). Based on this data, we investigated whether integrin β3 forms persistent nanoclusters or loosely held nanoclusters which frequently exchange molecules with the surroundings.

To investigate this in ∼100 nm regions within focal adhesions, we developed an analysis approach termed nanoKymo-FLAP (Fig. S1; performed on Photoactivatable Fluorescent proteins including PA-GFP, PA-RFP tagged to focal adhesion proteins), and nanoKymo-FRAP (Fig. S2; performed on fluorescent proteins GFP, mApple tagged to our protein of interest). To perform this analysis-a single fluorescence image of a live mouse embryonic fibroblast expressing a fluorescently tagged focal adhesion protein such as paxillin-mApple, spread on fibronectin-coated glass, is captured in Live-SR. In this image, a region of interest is marked along straight-lines with a single pixel width (activation region of interests, ROIs) across multiple adjacent focal adhesions (ROIs, Fig. S1a). For nanoKymo-FLAP, the same cell co-expresses a second photoactivatable fluorescent tagged adhesome protein e.g. vinculin-PAGFP. This was activated using the 405-nm laser in this predefined ROI and monitored over time using LiveSR (488-nm for PA-GFP, Fig. S1b). As shown in Fig. S1c, a kymograph was generated along the line ROIs and line profiles were created for high-density regions from the kymographs. These line profiles were then fitted using a one phase decay equation [22] to determine two parameters - (i) the half-time of residence of the mobile fraction within the focal adhesion and the (ii) percentage of immobile fraction (Ym). Once photoactivated, the dispersal of mobile fluorophores is expected to result in a reduction in fluorescent signal, while immobile fluorophores persist over time. Thus, a decrease in fluorescence intensity is indicative of the protein dynamics and mobility of the tagged protein in the activated ROI.

In contrast, in nanoKymo-FRAP, photobleaching of regular fluorescent proteins tagged to adhesome proteins was performed in similar ROIs as above (Fig. S2a) and monitored using LiveSR (Fig. S2b). Kymographs were generated along the line ROIs and line profiles of high intensity regions were created (Fig. S2c). These line profiles were fitted using an ‘exponential plateau’ equation [37, 47] to determine the half-time and mobile fraction. Once photobleached, the recovery of mobile fluorescent proteins is expected to result in an increase in fluorescent signal, while immobile fluorescent proteins do not recover. Thus, an increase in fluorescent protein intensity is indicative of increased protein dynamics and mobility in the bleached ROI. Since only sub-regions corresponding to a single line ROI were analysed, our nanoKymo-FLAP analysis allow us to probe sub-focal adhesion dynamics at close to the nanoscale dimension of similar to size of observed integrin nanoclusters within focal adhesions [1]. The confined illumination, combined with the LiveSR module provides a ∼100 nm resolution, close to the size of single nanoclusters [1]. These analyses were performed for multiple regions (at least 20) within focal adhesions for several cells (at least 5 cells from 3 repeats) to observe if distinct populations of a protein nanocluster with different dynamics are present within the focal adhesion.

Using nanoKymo-FLAP, we investigated the dynamics of integrin β3 using Mouse Embryonic Fibroblasts (MEFs) expressing integrin β3-PAGFP (PhotoActivatable GFP). We observed high and low-intensity regions with integrin β3 within focal adhesions (Fig. 1d, Movie S2.). Kymographs revealed that most of the high-intensity clustered regions persisted within the observation duration (5-10 mins) showing that the clustered regions were composed of a slow-exchanging or immobile population of integrins (Fig. 1e,f,g). Line profiles of high intensity regions from the kymographs were fitted to determine the immobile fraction, measured as a percentage of initial intensity upon photoactivation. In this manner, the dynamics of nanoregions corresponding to the nanocluster size within the focal adhesions, were determined.

Three analyses were performed on the dynamics data (i) <u>Ym Peak detection</u> - a histogram of all immobile fraction, Ym, measurements were plotted and the number of peaks was detected. This analysis allowed us to determine whether the population was uniform or consisted of distinct multiple peaks. This analysis detected two distinct peaks of integrin β3, one with a lower Ym which we termed - P_m_ with a peak at 41%, which implies that more than half the composition of these regions is mobile fraction, and another peak with a higher Ym termed P_im_ with a peak at 72% implying that these regions had higher fraction of immobile fraction (Fig. 1f). Hence, two types of regions are present in the focal adhesions, one with higher immobile fraction and one with lower immobile fraction. We observed that a focal adhesion does not consist of a single homogenous population of integrin dynamics, but two distinct populations are observed. We can’t ensure that we have precisely observed the dense nanoclusters of integrins, and some regions could be at the junction of nanoclusters and surrounding regions within the adhesions, similar to regions that overlap between water and island in an archipelago. However statistical analysis over multiple regions helped determine the distinct dynamics of nanoscale regions. (ii) <u>Ym Peak Fraction-</u> P_FWHM_-We analysed the relative fraction of total population in P_m_ & P_m_ gaussian fits. This analysis helped determine the relative population of each of the different peaks. We found that 0.30 of the population corresponded to the more mobile P_m_, while 0.27 corresponded to the higher immobile fraction P_im_ (Fig. 1f) and (iii) <u>Half-time analysis</u>, to be able to measure the dynamics of the mobile fraction. As expected, P_m_ had a 4.3x or longer half-life (13.37 s) within adhesions compared to P_im_ (3.09 s). The higher half-life of P_m_ would be because of the almost double mobile fraction (∼70%), which takes longer to exchange and reach a plateau, while P_im_ contains a lower mobile fraction (<30%), which would take a shorter time to exchange and reach a plateau, thus having a relatively shorter half-life. This result confirmed that, within the early focal adhesion, integrin β3 arranged in islands of immobile fraction in a sea of more mobile population (*P*=0.0013; Fig. 1g) [31, 48]. Taken together our results revealed that adhesions were composites of persistent regions with dense integrin β3 nanoclusters, as well as short-lived fast exchanging regions [31, 32, 46, 48]. Since activation of integrins promotes their immobilization, which is further stabilized by their connections to ECM and actin-binding proteins [31, 32], organisation of integrins into these long-lived immobile regions could provide a high spatial and temporal stability, which could function as robust platforms to organise signalling hubs of downstream signalling proteins within the focal adhesions.

### Nanoclusters of phosphorylated focal adhesion proteins are organized with ∼600 nm spacing

Due to integrin β3 being the early components of adhesions, we reason that integrin β3 clusters may serve to organise subsequently recruited adhesome components. First, we assayed if these proteins were organised as nanoclusters? To do so, we performed photoactivated light microscopy (PALM), that offers ∼25nm resolution using FAK-tdEos in FAK^−/−^ MEFs, or Vinculin-mEos2 in Vinculin^−/−^ MEFs, or Paxillin-mEos2 in MEFs respectively (Fig. S3). These also revealed the presence of clustered regions within focal adhesions similar to previous studies [46, 49–52].

To assess this, we next analysed the spatial organisation of key adhesome components with respect to integrin β3. We focussed on FAK as one of the primary kinases at the adhesions, paxillin as a LIM domain protein that translocates to the nucleus, and vinculin as a mechanosensitive adaptor protein. These 3 proteins could be recruited together [53]. Live-SR imaging of mApple-labelled FAK, Paxillin, or Vinculin revealed the nanoclustered organization of these proteins in focal adhesions (Fig. 2 a,b,c).

**Fig. 2.**
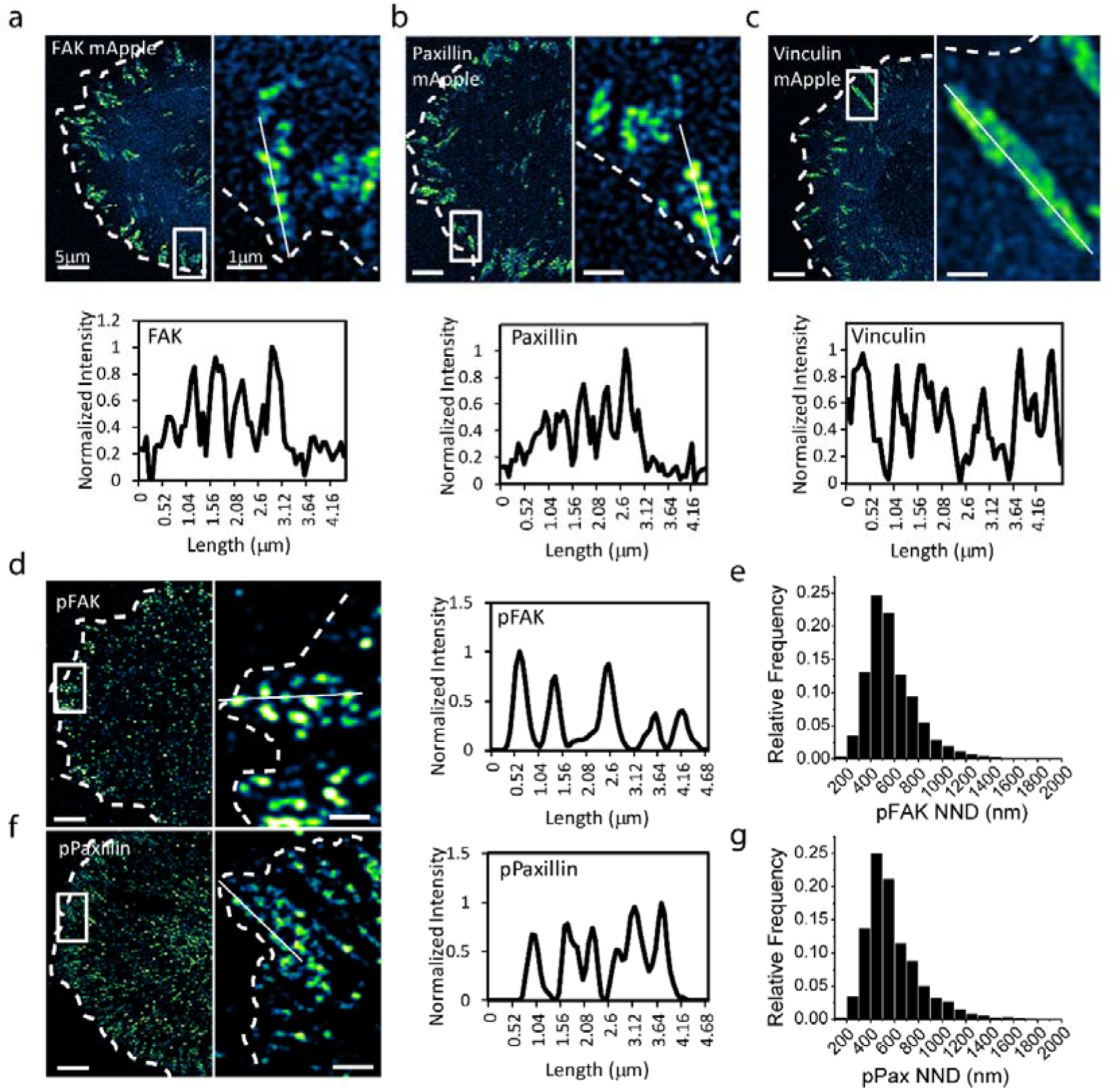
Integrin β3-associated focal adhesion proteins also organize in nanoclusters, with phosphorylated proteins at ∼600 nm distance. SR-confocal image of (a) FAK, (b) Paxillin, and (c) Vinculin along with zoom (right) of the white box. Line profile of white line showing normalized intensity peaks of nanoclusters on the bottom. (d) SR-confocal imaging of phospho-FAK (pFAK), along with zoom of the white box on the right. Line profile of pFAK along the while line in (d) showing nanoclusters spaced ∼600 nm. (e) Quantification of the nearest neighbour distance (NND) of phospho-FAK nanoclusters. N=31891 from 9 cells. (f) SR-confocal imaging of phospho-paxillin (pPax), along with zoom of the white box on the right. Line profile of pPax along the while line in (f) showing nanoclusters spaced ∼600 nm. (g) Quantification of the nearest neighbour distance (NND) of phospho-Pax nanoclusters. N=9520 from 11 cells. Scale bars for all left images, 5 µm. Scale bar for all zoom images (right), 1 µm.

Tyrosine phosphorylation events have been closely associated with focal adhesion signalling [54–58]. For instance, FAK phosphorylation at the Tyr397 residue leads to FAK activation [54, 55], while paxillin phosphorylation at Tyr31, Tyr118, Ser188, and Ser190 residues leads to activation of multiple signalling pathways [56–58]. Therefore, we next investigated the spatial arrangement of phosphorylated FAK (pFAK) and phosphorylated Paxillin (pPaxillin) using SR-confocal imaging. Surprisingly, while the spatial order in nanocluster organisation of total protein was not very clear, we observed that pFAK are organised into distinct nanoclusters. (Fig. 2d) The spatial organisation of these nanoclusters was quantified by nearest neighbour distance analysis that showed an average centre-to-centre distance of 580.85±1.27 nm (mean±S.E.M.; Fig. 2e). We obtained similar results with pPaxillin where approximately similar intervals (Fig. 2f) were determined, with an average centre-to-centre distance of 590.05±2.53 nm (mean±S.E.M.; Fig. 2g), close to the nearest neighbour distance of pFAK, also reported before [52]. These observations of very similar organisation of phosphorylated FAK and Paxillin compared to as integrin β3 (Fig. 1c) with a centre-to-centre distance of ∼600 nm suggests that β3 nanoclusters could be organising the nanoclusters of activated FAK i.e. phosophorylated FAK that subsequently phosphorylates other downstream proteins such as paxillin and organises them in clusters, hence could occur at and likely require the formation of nanoclusters of activated proteins. We analysed both these.

### Different focal adhesion proteins nanocluster in close proximity to integrin β3 nanoclusters

We sought to determine whether integrin β3 could spatial organisation of FAK, paxillin and vinculin in close proximity to its nanoclusters. We co-expressed integrin β3 along with either FAK-mApple, paxillin-BFP, or vinculin-mApple. Line profiles of Live-SR images demonstrated that nanoclusters of all the three proteins FAK, paxillin, and vinculin can be found in close proximity to integrin β3, with high overlap (Fig. 3a,b,c). We further performed FLAP by co-photoactivating integrin β3-PAGFP along with Vinculin-PATagRFP, while using Paxillin-mIFP (far red fluorescent protein) to mark ROIs and monitor adhesions (Figure 3d). Examination of fluorescence intensities over time (Fig. 3d), revealed that both paxillin and vinculin clusters closely followed integrin β3 clusters. Initially, most nanoclusters were observed to coincided, suggesting that integrin β3 nanoclusters could serve to nucleate nanocluster of different focal adhesion proteins. However, vinculin clusters started to disappear first, followed by paxillin, even when integrin β3 clusters persist (Fig. 3d). These suggest that the paxillin and vinculin nanoclusters were more dynamic compared to integrin β3 nanoclusters. This could be due to the fact that these molecules carry signals elsewhere within the cell for further downstream signalling or other functions. For instance, activated Paxillin, as a LIM domain protein, translocates to the nucleus to regulate gene expression in response to mechanical force [18]. Hence upon activation at the focal adhesion it would leave the focal adhesions, to carry the signal to the nucleus. Similarly, vinculin that is located further away from the integrins, in the layers of focal adhesion [40], has an actin-organising function [59]. Thus, this would imply that these protein nanoclusters have different dynamics within the focal adhesion compared to integrin.

**Fig. 3.**
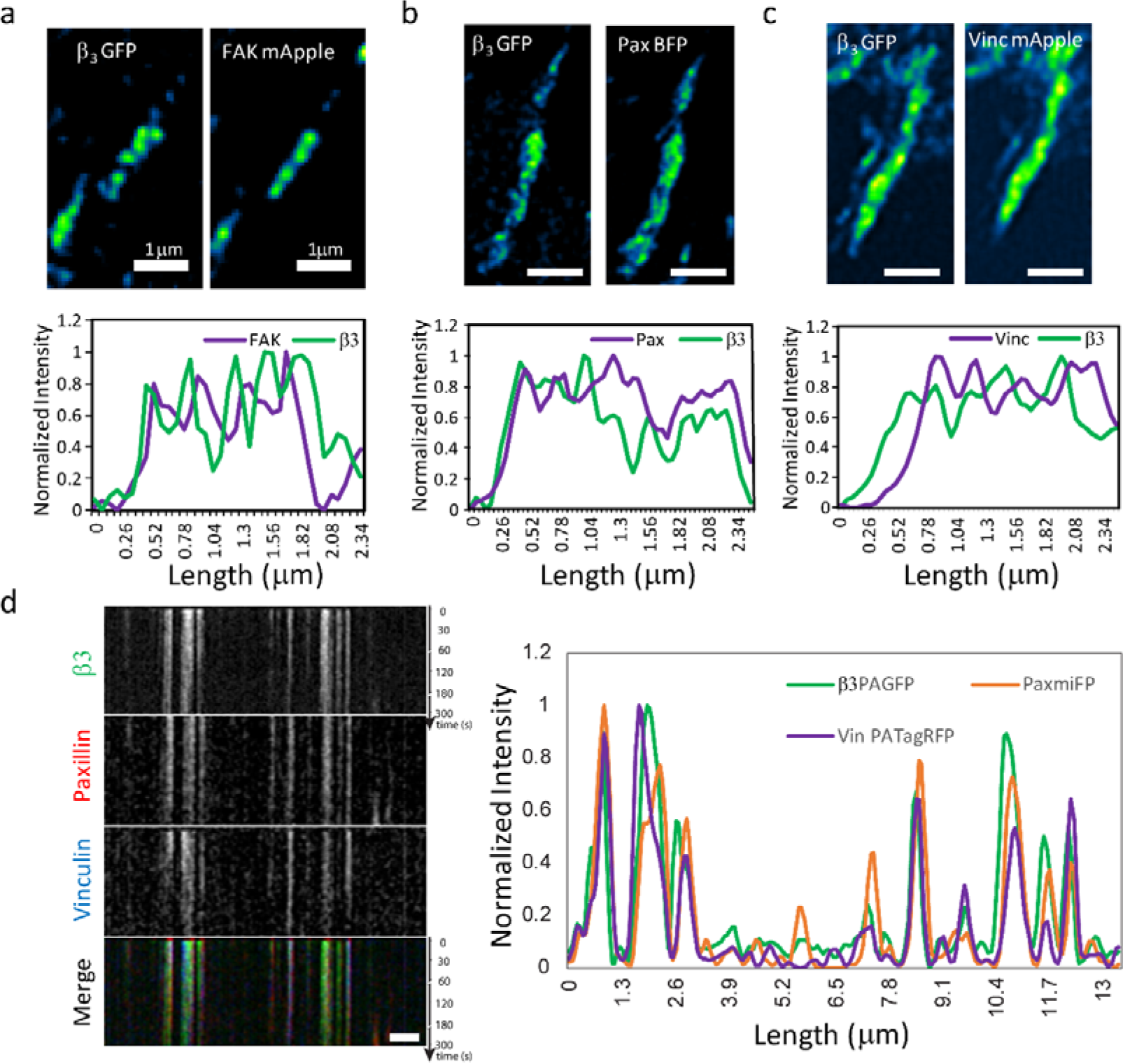
Different focal adhesion proteins nanocluster in close proximity to integrin β3 nanoclusters. SR-confocal images of MEFs co-overexpressing integrin β3-GFP with (a) FAK-mApple, (b) Paxillin-BFP, and (c) Vinculin-mApple. Line profile showing normalized intensity peaks of nanoclusters on the bottom. Scale bars for a-c, 1 µm. (d) Kymograph of Fluorescence Loss after Photoactivation (FLAP) of MEFs co-overexpressing integrin β3-PAGFP, Vinculin-PA-TagRFP, and Paxillin-mIFP, along with the line profiles at activation time on the right. Scale bar for d, 2 µm.

### NanoKymo-FLAP/FRAP reveals differential dynamics of core focal adhesion proteins

To further investigate the dynamics of focal adhesion proteins nanoclusters, we performed nanoKymo-FLAP/FRAP analysis of FAK, vinculin and paxillin.

FAK-PATagRFP was used to perform nanoKymo-FLAP analysis. Ym histogram of the immobile fraction demonstrated two distinct populations, one reaching a plateau at ∼35%, and the other reaching ∼70% of the initial intensity, comparable to integrin β3 (Fig. 4a, Movie S3.). Ym Peak detection analysis showed that FAK clusters can be categorised by two distributions, with peaks at 36±10% (mean±s.d.; comparable to P_m_) and 71±24% (mean±s.d.; similar to P_im_; Fig. 4b). Ym Peak Fraction showed that 0.375 of the population belonged to P_m_, while 0.19 of the population belonged to P_im_. Half-time analysis showed that both these populations differed significantly, with the half-time of P_m_ being 10.4±1.5 s (mean±S.E.M.) and the half-time of P_im_ being 3.1±0.6 s (mean±S.E.M.; *P*=0.0041; Fig. 4c), the reasoning of P_im_ having a shorter half-time is similar to that of integrin, where far fewer molecules exchange so the population reaches a plateau sooner. To further ascertain the nanoKymo-FLAP results, we also performed nanoKymo-FRAP analysis of FAK-mApple. This also revealed two distinct populations with similar plateau values, one reaching a plateau at ∼40%, and the other reaching ∼60% of the initial intensity (Fig. S4a, Movie S4.). Ym Peak detection analysis showed that FAK clusters had two populations, with recovered fractions of 39±14% (mean±s.d.; P_im_) and 65±25% (mean±s.d.; P_m_; Fig. S4b). Half-times of both these populations differed by 2 times, with the half-time of P_im_ being 6.1±0.7 s (mean±S.E.M.) and the half-time of P_m_ being 12.9±1.8 s (mean±S.E.M.; *P*=0.0061; Fig. S4c). The complementary Gaussian peaks obtained for P_m_ and P_im_ also highlight the complementary nature of FRAP and FLAP measurements. Moreover, FAK is similarly distributed to integrins. It’s not possible to tag pFAK in real time, but most likely the larger immobile fraction would contain pFAK, we test this hypothesis subsequently using FAK mutants.

**Fig. 4.**
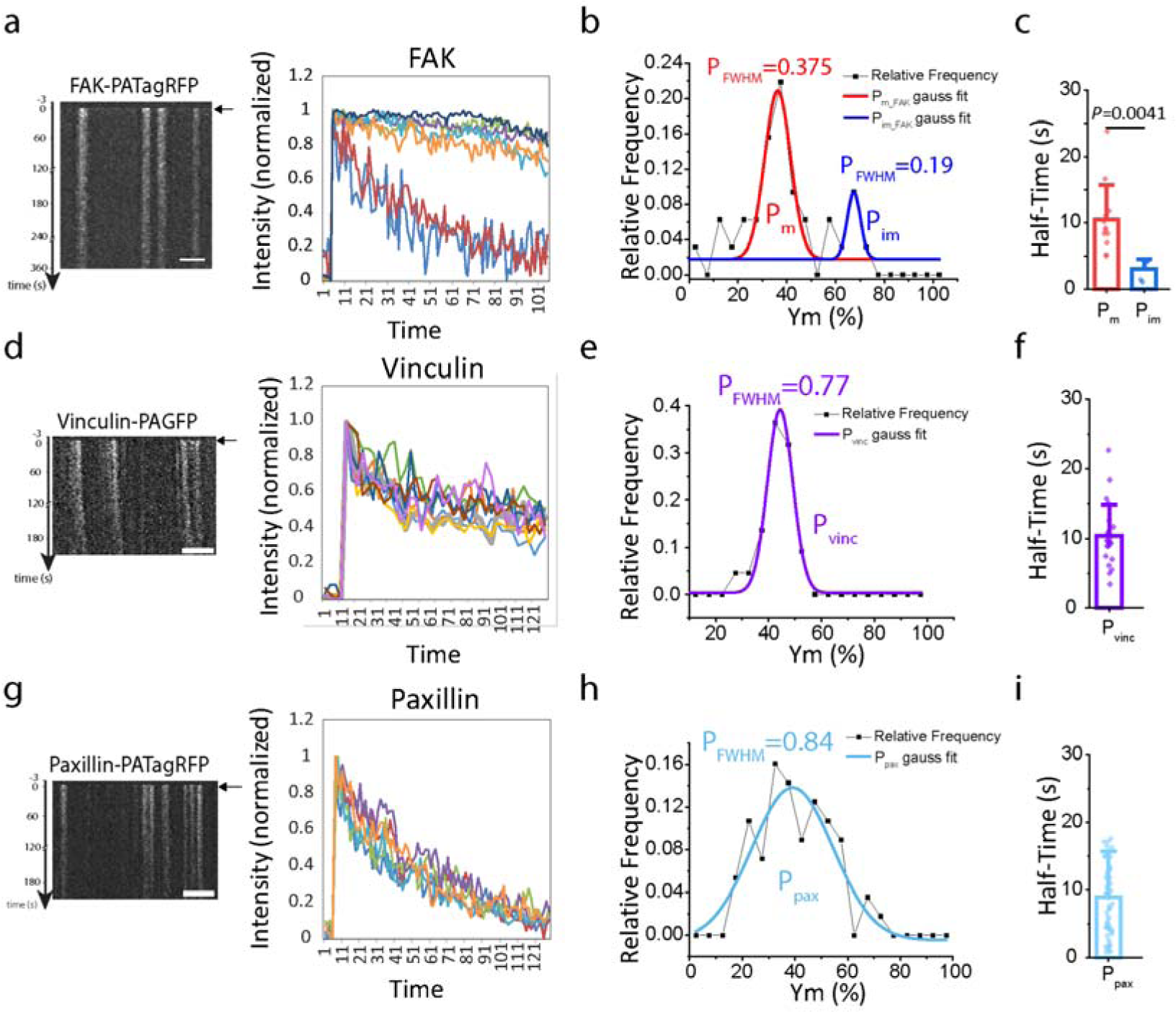
Differential nanocluster dynamics of focal adhesion proteins. (a) Kymograph of nanoKymo-FLAP for FAK-PATagRFP overexpressing MEFs along with the representative normalized intensity vs time graphs for nanoKymo-FLAP. (b) Relative frequency (histogram) of Ym calculated from fitting intensity vs time plots, along with Gaussian fits of the 2 populations. N=32 from 5 cells. (c) Mean±S.D. plot of half-time of fluorescence loss of the 2 populations, along with the individual points. N=12 for P_m_, N=6 for P_im_. (d) Kymograph of nanoKymo-FLAP for Vinculin-PAGFP overexpressing MEFs along with the representative normalized intensity vs time graphs for nanoKymo-FLAP. (e) Relative frequency (histogram) of Ym calculated from fitting intensity vs time plots, along with Gaussian fit of the population. N=22 from 5 cells. (f) Mean±S.D. plot of half-time of fluorescence loss of the population, along with the individual points. (g) Kymograph of nanoKymo-FLAP for Paxillin-PATagRFP overexpressing MEFs along with the representative normalized intensity vs time graphs for nanoKymo-FLAP. (h) Relative frequency (histogram) of Ym calculated from fitting intensity vs time plots, along with Gaussian fit of the population. N=56 from 7 cells. (i) Mean±S.D. plot of half-time of fluorescence loss of the population, along with the individual points. P_FWHM_ denotes the fraction of population lying in the full width half maxima of the Gaussian fit. *P* value was evaluated by two-tailed t-test.

We analysed other adhesome proteins. In contrast to FAK, Ym Peak detection analysis of nanoKymo-FLAP of vinculin and paxillin revealed a single population. nanoKymo-FLAP analysis of Vinculin-PAGFP demonstrated a comparable loss of fluorescence intensity, reaching a plateau at ∼44% of the initial intensity (Fig. 4d, Movie S5.). Vinculin nanocluster dynamics was also characterised by a single population, with a peak at 44±5% (mean±s.d.; Fig. 4e) and a half-time of 10.1±0.9 s (mean±S.E.M.; Fig. 4f). Similarly, nanoKymo-FLAP analysis of Paxillin-PATagRFP revealed a rapid loss of fluorescence intensity following photoactivation, reaching a plateau at ∼40% of the initial intensity (Fig. 4g, Movie S6.). Ym Peak detection analysis revealed that Paxillin clusters exhibited a single population, with a peak at 39±15% (mean±s.d.; Fig. 4h) and a half-time of 8.9±0.9 s (mean±S.E.M.; Fig. 4i).

These data showed that protein clusters in focal adhesions have different dynamics, similar to β3, FAK clusters were more distinctly separated into mobile (high half time) and immobile clusters (low half-time) as observed by both nanoKymo-FLAP and nanoKymo-FRAP. These results imply that nanoclusters of FAK, a key signalling kinase that is immediately downstream of integrin and phosphorylates other proteins at the adhesions, are spatially and temporally correlated with integrin nanoclusters. In contrast, nanoclusters of vinculin and paxillin, potentially activated by FAK [60, 61]; both potentially bring about function elsewhere in the cell and exhibit similar single population dynamics (mixed mobile and immobile fractions) with a comparable half-time of ∼10 s. The relative decrease in FAK half-time as compared to Paxillin and Vinculin is consistent with previous literature [48], albeit with differences in values. This difference could be caused by multiple reason including, the size of ROI within the focal adhesions, or due to different phases of cell spreading being analysed, whereby in this study we focused on the early spreading phase (15-50 mins), before the recruitment of integrin β1 whereas Stutchbury et al [48] focus on the late spreading phase after overnight spreading.

Importantly, this result shows that FAK, that is phosphorylated directly by force-dependent integrin signalling, and in turn activates other proteins [34, 62, 63], does so by forming long-lived nanoclusters close to integrin nanoclusters to facilitate biochemical signalling in response to integrin activation. This FAK mediated biochemical signalling is translated to downstream signalling events such as, phosphorylation and activation of LIM domain protein, paxillin, which then, can transmit the biochemical signal to the nucleus and activates downstream gene expression [18]. This demonstrates that long lived FAK nanoclusters are organized by integrin nanoclusters. Can they function as signalling hubs?

### NanoKymo-FLAP/FRAP analysis reveals the dependence of immobile clusters on FAK phosphorylation

We hypothesised that immobile FAK nanoclusters are phosphorylated and can function as signalling hubs. Since FAK activation was known to affect focal adhesion stability [60, 64, 65], we first inhibited FAK activity using PF-562271 and performed nanoKymo-FRAP analysis to assess FAK cluster dynamics. Interestingly, in the Ym Peak detection analysis, instead of two distinct populations, we observed a single population with a recovered fraction of 52±12% (mean+s.d.; Fig. 5a. Movie S7.), and a half-time of 11.77±1.24 s (mean±S.E.M.), similar to the P_m_ population in untreated FAK (Fig. 5b,c, *P*=0.6032). The loss of P_im_ population upon FAK inhibition thus suggests that the stability of immobile fraction of FAK is dependent on FAK activation. We further expressed FAK-Y397F, the non-phosphorylatable, and hence inactive mutant of FAK [66] in FAK^−/−^ cells. In Ym Peak detection analysis of nanoKymo-FLAP, FAK-Y387F mutant revealed the loss of the two distinct peaks, with a single peak corresponding to P_m_, containing larger mobile fraction (0.83) with a peak at 30±14% (mean+s.d.; Fig. 5d, e, Movie S8.), and a half-time of 11.21±1.33 s (mean±S.E.M.; Fig. 5f), similar to the P_m_ nanoKymo-FLAP population in untreated FAK (*P*=0.7285). These findings thus indicated that the phosphorylation of FAK at Tyr397 is necessary for stable immobile population of FAK, and that stable immobile pFAK is the signalling hub for downstream signalling.

**Fig. 5.**
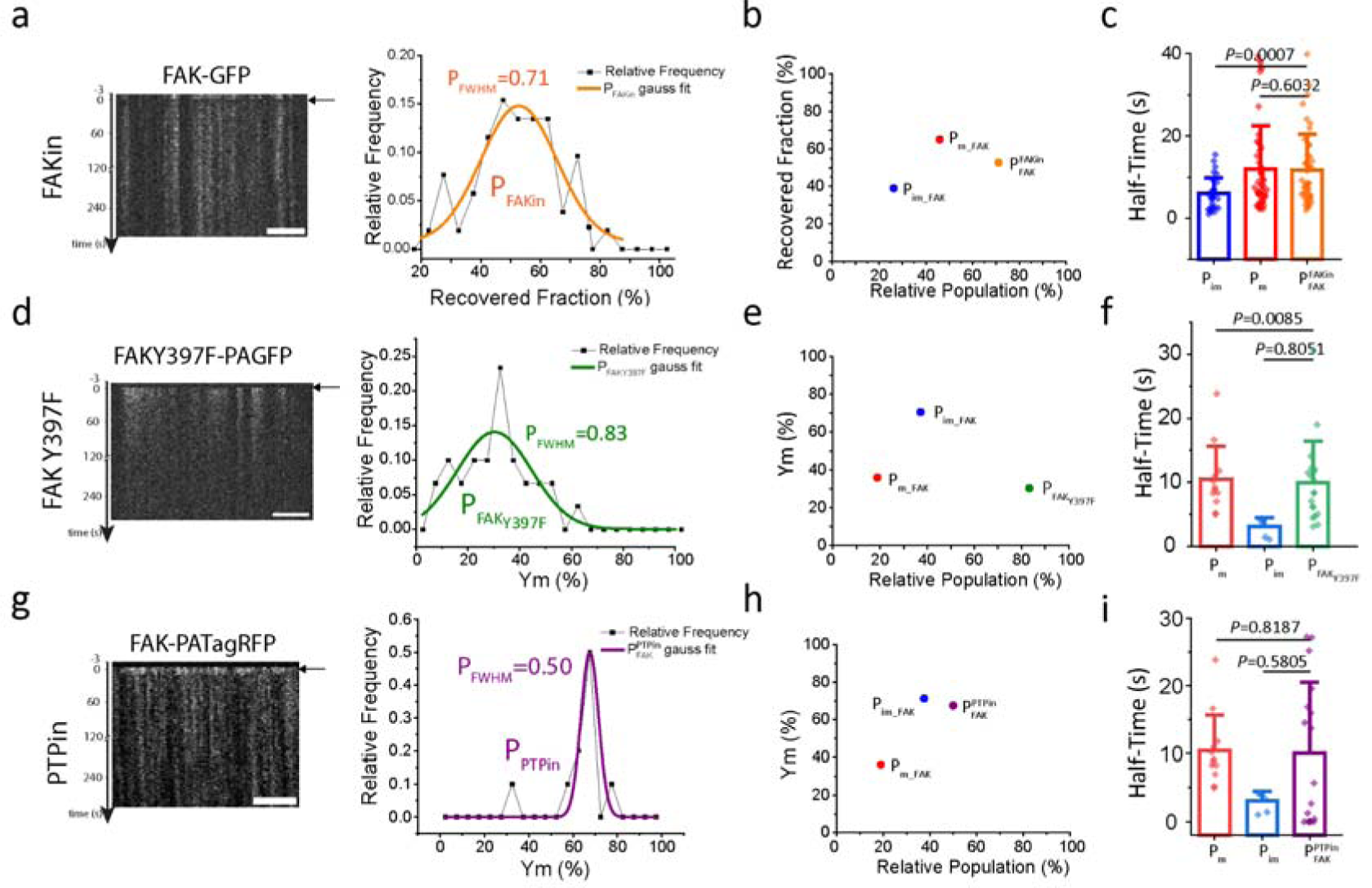
Presence of immobile clusters requires phosphorylation of FAK. (a) Kymograph of nanoKymo-FRAP for FAK-GFP for MEFs treated with 10 µM PF-562271 (FAKin) along with the histogram of recovered fraction. The distribution was calculated from fitting intensity vs time plots and plotted along with the Gaussian fit of the population. N=43 from 8 cells. (b) Recovered nanoKymo-FRAP fraction vs Relative population plot comparing the FAKin population with control FAK nanoKymo-FRAP populations. (c) Mean±S.D. plot of half-time of nanoKymo-FRAP control vs FAKin populations. (d) Kymograph of nanoKymo-FLAP for FAKY397F-PAGFP overexpressing MEFs along with histogram immobile nanoKymo-FLAP population and Gaussian fit of the population. N=27 from 5 cells. (e) Immobile nanoKymo-FLAP fraction vs Relative population plot comparing the FAKY397F population with control FAK nanoKymo-FLAP populations. (f) Mean±S.D. plot of half-time of nanoKymo-FLAP control vs FAKY397F populations. (g) Kymograph of nanoKymo-FLAP for FAK-PATagRFP for MEFs treated with 2.5 µM PTP inhibitor (PTPin) along with the histogram of recovered fraction and Gaussian fit of the population. N=20 from 5 cells. (h) Immobile nanoKymo-FLAP fraction vs Relative population plot comparing the PTPin population with control FAK nanoKymo-FLAP populations. (i) Mean±S.D. plot of half-time of nanoKymo-FLAP control vs PTPin populations. P_FWHM_ denotes the fraction of population lying in the full width half maxima of the Gaussian fit. P values were evaluated by Mann-Whitney tests.

Since tyrosine phosphorylation is counter-balanced by tyrosine dephosphorylation in cells [67, 68], we hypothesised that treatment with the appropriate PTP inhibitor would lead to increased FAK activation, leading to a higher immobile fraction. Force dependent protein tyrosine phosphatase PTPN12 is known to dephosphorylate and thus deactivate FAK [67, 68]. So, upon inhibition of PTPN12, we would expect loss of P_m_ and increase in P_im_ population in the histogram analysis of nanoKymo-FLAP. To inhibit PTPN12, we treated MEFs overexpressing FAK-PATagRFP with PTPN12 inhibitor and performed nanoKymo-FLAP analysis. Indeed, Ym Peak detection analysis revealed that this treatment resulted in a single population of FAK, with a peak at 67±4%, similar to the P_im_ nanoKymo-FLAP population in untreated FAK (mean±s.d.; Fig. 5g, h, Movie S9.), and a majority of population in half-time close to the P_im_ population in untreated FAK (16.49±3.48 s, mean±S.E.M., Fig. 5i). This demonstrates that the 2 distinct populations of FAK where P_im_ corresponds to nanoclusters of immobilised, signalling active, phosphorylated FAK and P_m_ to unphosphorylated and relatively short-lived single molecules of FAK within the focal adhesion [69]. Taken together, our results showed that signalling active phosphorylated FAK immobilises in nanoclusters that serve as signalling hubs.

### Stability of Integrin β3 nanoclusters is dependent on contractility

FAK is a force dependent kinase, which is phosphorylated and activated on stiff substrates [14, 16, 70]. Hence, if the stable nanoclusters of FAK are signalling hubs necessary for signalling, absence of force would eliminate these nanoclusters. Therefore, we sought to determine whether the stability of the nanoclusters is contractility dependent. We treated cells with ROCK inhibitor Y-27632, which is known to reduce contractility, drastically reduce adhesion formation, focal adhesion lifetime, and increase focal adhesion disassembly rate [20–23]. Upon Y-27632 treatment after activation, we observed that integrin β3-PAGFP nanoclusters disassembled rapidly within seconds (Fig. 6a,b, Movie S10.), suggesting that reduction of contractility results in low immobile fraction (Fig. 6a,b). nanoKymo-FLAP Ym Peak detection analysis revealed that Y-27632 treated cells exhibited a Ym of 25±8% (mean±s.d.; Fig. 6c,d) and a high half time of 20.5±3.7 s (Fig. 6e), similar to the P_m_ population in wild-type. This result suggests that the presence of actomyosin contractility is required for nanocluster stability.

**Fig. 6.**
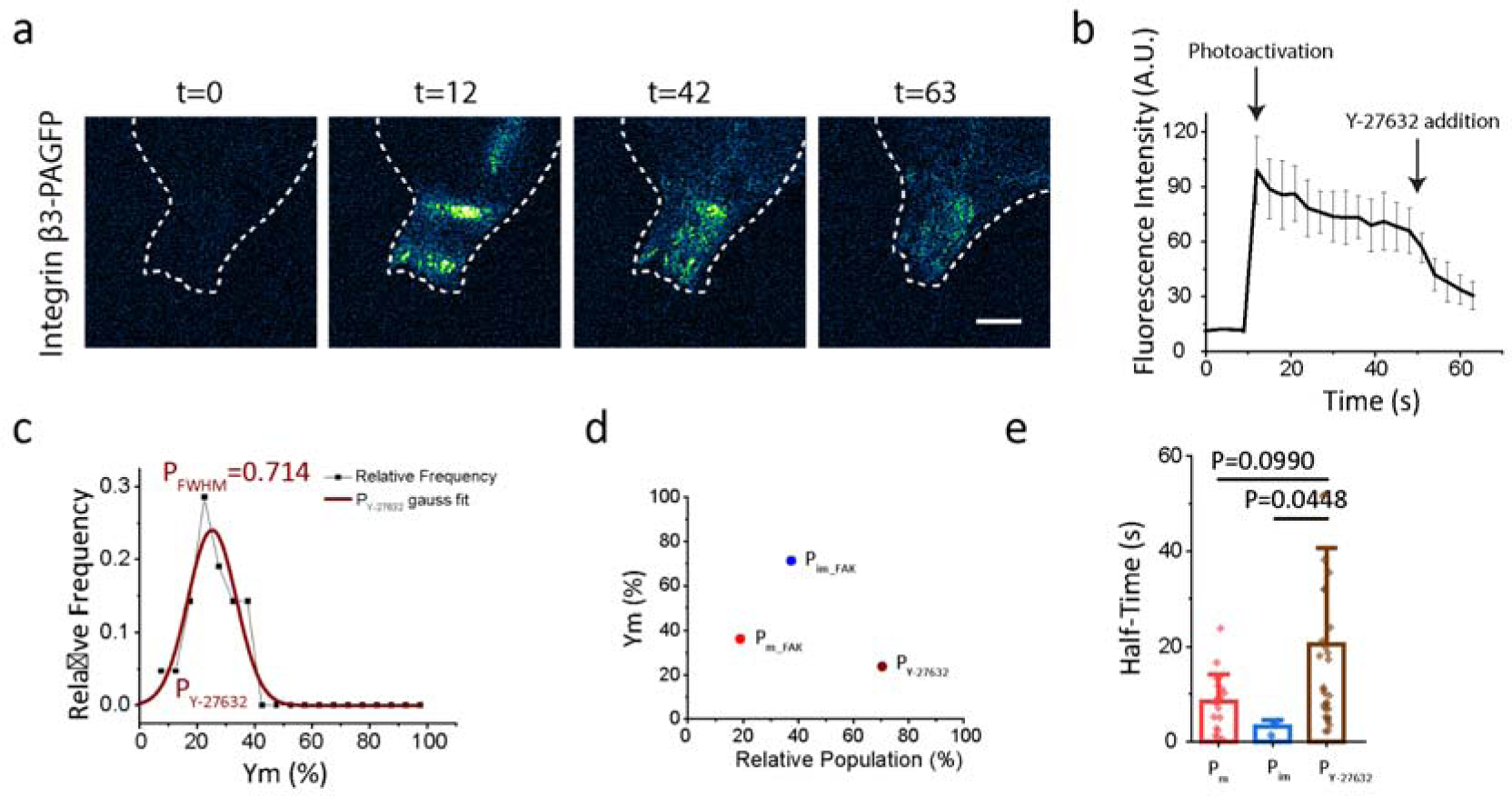
Immobilization of Integrin β3 nanoclusters is force-dependent. (a) nanoKymo-FLAP imaging of integrin β3 activated at t=12 s and treated with Y-27632 at t=42 s. Dashed line represents cell boundary. Scale bar, 5 µm. (b) Fluorescence intensity profile showing photoactivation, and Y-27632 addition time points. (c) Relative frequency of stable integrin β3 nanoKymo-FLAP population for MEFs overexpressing integrin β3-PAGFP and treated with 10 µm Y-27632 along with the Gaussian fit of the population. N=21 from 5 cells. (d) Stable nanoKymo-FLAP fraction vs Relative population plot comparing the Y-27632 population with control populations. (e) Mean±S.D. plot of half-time of nanoKymo-FLAP control vs Y-27632 treated populations. PFWHM denotes the fraction of population lying in the full width half maxima of the Gaussian fit. P values were evaluated by two-tailed t-test.

### Tyrosine phosphorylation stabilizes immobile clusters regardless of contractility

PTPN12 is a force dependent phosphatase that dephosphorylates FAK in absence of force [67, 68]. Hence it would be required for elimination of FAK signal by dephosphorylation. Therefore, we hypothesized that if PTPN12 is inhibited and followed by elimination of traction force (by treatment with Y-27632), then FAK cluster disassembly could be inhibited as pFAK would still persist. Upon treatment with PTPN12 inhibitor, we observe that it led to larger focal adhesions (Fig 7e), with a 27% increased total adhesion area as compared to untreated cells (Fig. 7a,e, Fig 5g). In contrast, treatment with Y-27632 led to rapid disassembly of focal adhesions, cell spreading out and formation of small nascent adhesions (Fig 6, 7a, 7e). Adhesions present before treatment with Y27632 did not persist, as observed in the merge (Fig 7a, middle panel). in addition, we observed that focal adhesions became unstable, disassembled rapidly, and the cells had a 15% lower total adhesion area (Fig. 7a,e,f, Fig. 6). However, when the cells were pretreated with PTPN12 inhibitor for 30 mins, followed by an elimination of traction force by treatment with ROCK inhibitor, Y-27632 (15 minutes), we observed that the cell spread out, several of the adhesions present prior to the treatment with Y-27632, continued to persist even throughout the treatment (Figure 7a, lower panel, merge) indicating that adhesions did not disassemble rapidly. Next, dynamics of FAK was measured using nanoKymo-FLAP. Ym peak analysis of FAK-PAGFP showed an increase in the Ym peak value (42±12%; Fig. 7b,c) upon treatment with both inhibitors (as described above) as compared to treatment with only Y-27632 (25±8%, Figure 7b, c). The half-time of the population also decreases from 20.5±3.7 s to 7.6±0.9 s (*P*=0.0012; Fig. 7d), respectively. The increased Ym peak value and decreased half-time value, suggested that when PTPN12 activity is reduced, the immobile population of FAK increases even upon reduction of traction force. This increase in the immobile fraction, indicates that signal disassembly required dephhosphorylation, and that phosphorylated FAK nanoclusters do not disassemble without the presence of PTPN12 activity. Pre-treatment with PTPN12 inhibitor also led to an increase in total adhesion area normalised to cell area (Fig. 7e, f), and instead of disassembling, a significant portion of these adhesions persisted even in the absence of traction force, indicating that phosphorylation is a key signal in maintaining persistent signalling hubs, and dephosphorylation is required for disassembly of the immobile nanocluster signalling hubs for removal of the signal (Fig. 5).

**Fig. 7.**
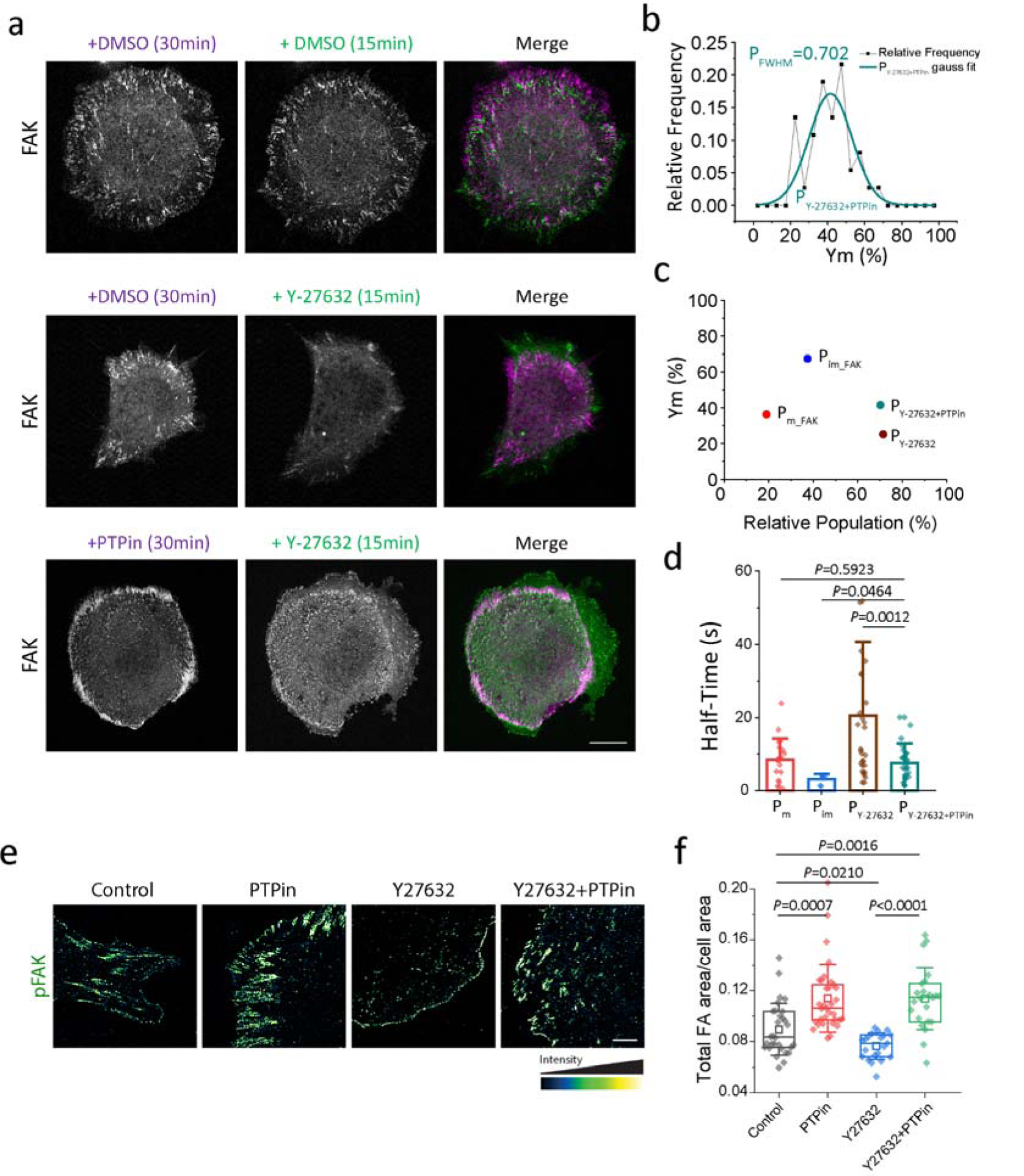
Phosphorylation can rescue presence of immobile clusters in the absence of force. (a) Fluorescence images of MEFs over-expressing FAK-mApple, treated with PTP inhibitor (PTPin) or DMSO for 30 min and Y-27632 or DMSO for another 15 mins, along with the merge. Dashed line represents cell boundary. N=44 from 6 cells. Scale bar, 10 µm. (b) Relative frequency of stable FAK nanoKymo-FLAP population for MEFs overexpressing FAK-PAGFP and treated with 10 µM Y-27632 with PTPin treatment along with the Gaussian fit of the population. P_FWHM_ denotes the fraction of population lying in the full width half maxima of the Gaussian fit. N=37 from 5 cells. (c) Stable nanoKymo-FLAP fraction vs Relative population plot comparing the Y-276329+PTPin population with control and Y-27632 populations. (d) Mean±S.D. plot of half-time of nanoKymo-FLAP control vs Y-27632 vs Y-27632+PTPin treated populations. (e) Control, PTPin treated, Y-27632 treated, and Y-27632 and PTPin treated cells spread and immunostained for FAK. Dashed line represents cell boundary. Scale bar, 5 µm. (f) Quantification of total adhesion area normalized to the cell area. Box plots with data overlay display the upper and lower quartiles and a median, the circle represents the mean, and the whiskers denote the standard deviation values, along with the individual points and P values. P values were evaluated by one-way ANOVA and Tukey’s post hoc test.

### Multiplexing nanoKymo-FRAP with nanopatterning reveals the dependence of nanocluster stability on nanocluster spatial organization

The observation of the ∼600 nm characteristic distance between phosphorylated nanoclusters of FAK and Paxillin is similar to the previously shown ∼600 nm optimal intercluster distance between ligand presentation that is required for optimal cell spreading and adhesion maturation [41]. Further, if this distance is halved to ∼300nm or almost doubled to 1000nm, adhesions do not form, and cells do not spread [41]. This intercluster distance of 600 nm dictates integrin nanocluster formation, which could dictate the directs formation of long lived immobile FAK nanoclusters, which are key signalling hubs. To present ligands at varying predetermined intercluster distances to the cell, we presented cyclo-RGD, an integrin binding ligand, on hexagonal array of nanodisc with a diameter of 100 nm separated by an interdisc distance of 300 nm, 600 nm or 1000 nm. To achieve this, we nanopatterned Ti discs of 100nm separated by the required distance in a hexagonal disc pattern using electron-beam lithography (EBL). This Ti pattern was oxidised to TiO2 and functionalized with cyclo-RGD using biotin-labelled phophoinositol bis-phosphate, which was in turn bound to fluorescently labelled neutravidin as the linker between the TiO2 and cyclo-RGD [41]. The advantage of using nanopatterned substrates was that it provided an important parameter where we can measure the dynamics of clusters directly on precisely spaced ligands, which was not possible on continuous substrates. In addition, unlike Au, TiO2 patterns allows us to image dynamics of molecules at the focal adhesions. Hence this offers an additional functional scenario to measure FAK dynamics when focal adhesions form versus do not form.

When MEFs overexpressing mApple-FAK were spread on 600nm, optimally spaced TiO_2_ nanopatterned substrates we observed that a majority of the FAK nanoclusters coincided with the nanopatterned ligands, further implying that activated integrin nanoclusters seed signalling nanoclusters of FAK (Fig. 8a,b, Movie S12., [71]). Further, this geometry, allows us to precisely define where the integrin nanoclusters are and only measure the dynamics of FAK on top of the integrin clusters seeded by the ligand presenting nanodisc. nanoKymo-FRAP was performed and FAK dynamics was measured precisely on the nanodiscs. On 600 nm spacing, FAK demonstrated a single recovered population, with a peak at 25±13% (mean±s.d.; Fig. 8b,c), suggesting the presence of stable nanoclusters on the immobile ligand nanodiscs. To investigate whether altering the intercluster spacing has an effect on the FAK nanocluster dynamics, when the spacing was decreased to half the optimal spacing value i.e. 300 nm, or to almost double i.e. 1000 nm using nanopatterned substrates (Fig. 8d), nanoKymo-FRAP analysis of FAK revealed a single recovered population with a peak at 50±22% on 300nm spaced nanodiscs(mean±s.d.; Fig. 8d,e,f, Movie S12.), showing reduced stability and increased mobile fraction of FAK nanoclusters.. Further, nanoKymo-FRAP analysis of FAK in cells spread on the 1000 nm intercluster distance revealed two distinct recovered populations, with peaks at 31±5% (mean±s.d.), and 59±10% (mean±s.d.; Fig. 8d,e,f, Movie S12.), showing an increased mobile population when the intercluster distance is increased to 1000 nm. Peak Fractions showed that 0.375 of the population belonged to P_im_, while 0.44 of the population belonged to P_m_ indicating an increase in mobile fraction present on the nanodiscs Taken together, these results demonstrate that nanoclusters arranged with an intercluster distance of 600 nm had a higher stability as compared to the nanoclusters formed with an intercluster distance of 300 nm or 1000 nm, indicating that persistent FAK nanoclusters are formed only when the external geometry is conducive.

**Fig. 8.**
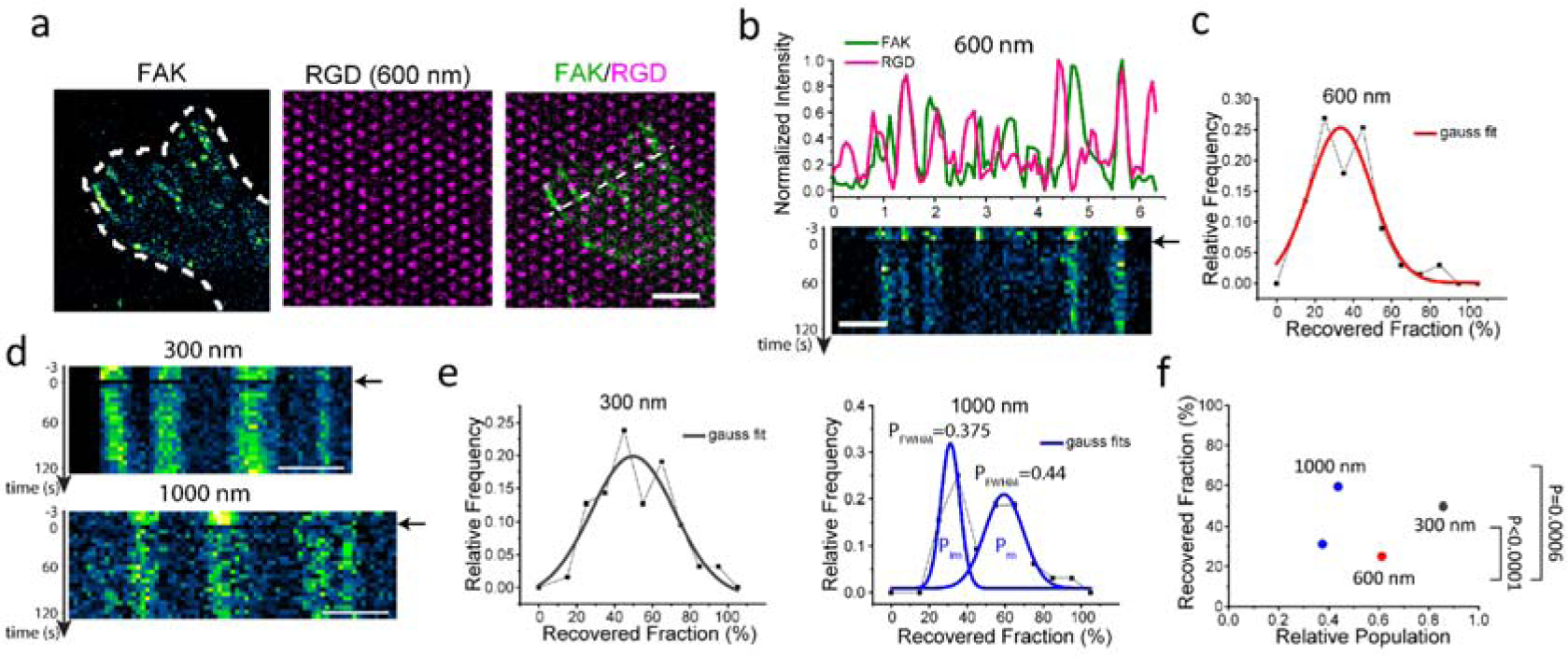
Nanocluster spacing determines cluster stability. a) Representative super-resolution confocal image of MEFs overexpressing FAK-mApple spread on nanodisc substrates with 600 nm intercluster distance, along with merge. The nanodiscs were labelled with Neutravidin-Dylight 650 to mark RGD. The dashed line represents cell boundary. Scale bar, 2 µm. (b) Kymograph of FAK along the dotted white line in (a) along with the intensity line profile of the first frame for FAK and RGD on 600 nm intercluster distance. Black arrow denotes timepoint when FRAP was performed. Scale bar, 1 µm. (c) Relative frequency of recovered fraction calculated from fitting intensity vs time plots of FAK for intercluster distances of 600 nm along with Gaussian fits of the populations. N=67 from 8 cells for 600 nm. (d) NanoKymo-FRAP kymographs of FAK on nanodisc patterns of 300 and 1000 nm intercluster distances. Black arrow denotes timepoint when FRAP was performed. Scale bars, 1 µm. (e) Recovered fraction vs Relative population plot for 300 nm and 1000 nm intercluster distances, along with the Gaussian fits. N=63 from 8 cells for 300 nm, N=32 from 6 cells for 1000 nm. (f) Recovered nanoKymo-FRAP fraction vs Relative population plot comparing the different intercluster distances.

## Discussion

Nanocluster assembly is a widely observed phenomenon in multiple transmembrane receptors; how these clusters play a role in signalling, is poorly understood [1–7]. Since signalling by single receptor can lead to false positives and poor control in the signalling process, nanoclusters can help serve as platforms for robust signalling [31, 72, 73] and provide a threshold (or critical minima) for activating specific signalling. Nanoclustering of various transmembrane receptors can form via mechanisms such as protein-protein interaction leading to phase separation [72, 74, 75], lipid interactions [76–79], reorganization of actin cytoskeleton [2, 80–82]. It is hypothesized that nanoclusters can provide various advantages such as increase in avidity, and help in maintaining active signalling [8]. Here, we investigate this hypothesis and define key properties of nanoclusters which allows it to function as a signalling hub by understanding nanoclusters of transmembrane receptors, integrin. Focal adhesions comprise of nanoclusters of integrin and other adhesome proteins [1, 30, 46, 49–52]. However, are these nanoclusters constantly cycling or are they stable over time is not well understood. Single molecule imaging has revealed that individual integrin molecules can be either immobile or mobile [31], however are there multi-molecular assemblies, perhaps as signalling hubs in form of nanoclusters, and do they bring about signalling? The behaviour of integrins within the adhesions at the scale of the nanocluster, i.e. ∼100nm [1] as an assembly of several molecules is not understood. Numerous previous studies have focused on the regulation focal adhesion dynamics either at the micron-scale, by tracking full focal adhesions [33, 34] or using FRAP [35, 36], or at the single-molecule level, using single-particle tracking [31, 46], however studies focusing on the dynamics of nanoclusters have been elusive. To study the nanocluster dynamics, we used nanokymo-FLAP/nanokymo-FRAP to activate/bleach thin lines of ROIs across focal adhesions, wherein thin line ROIs are activated and tracked within a focal adhesion. Since not all regions would be nanoclusters, hence we see if within these thin strips ROIs there is a percentage of stable/immobile versus unstable/ mobile regions. The stable immobile regions potentially contain signalling nanoclusters or hubs. The proportion of mobile versus immobile nanocluster regions was measured and each of their half-life was calculated. This was measured for different focal adhesion proteins which serve distinct functions-integrin transmembrane receptors bridge the ECM to the cell interior, that upon activation activates FAK [13–16, 83], a key force-dependent kinase required to activate downstream signalling molecules. For instance paxillin which is a LIM domain protein that upon force-dependent phosphorylation is translocated to the nucleus to bring about regulation of specific gene expression [18, 19]. This measurement was used as a benchmark where dynamics of nanoclusters in the wild type cells were compared to mutant or drug treated cells to understand if dynamics is related to function, which is hypothesized to be brought about by nanoclusters.

Firstly, nanoclusters, not single molecules of integrins, act as a signalling hub. Using nanokymo-FRAP and nanokymo-FLAP, we observed that integrin nanoclusters are distinctly immobile, where a nanocluster persist over time. This shows a multi-molecular assembly (as a nanocluster) persisting over time. Exchange of individual molecules would lead to changes in fluorescence intensity within the nanocluster, which we do not observe. This indicates that nanoclusters, not individual single molecules, are stable over time, satisfying the first criterion of it being a signalling hub. This stability of a nanocluster as a signalling hub serves multiple functions: it improves signal to noise ratio, whereby signalling via single molecules can create false positives, whereas if it is orchestrated by a group of integrins it increases the local concentration of a signal which would be a positive signal only above a threshold thereby having a robust signalling outcome [8, 12].

Secondly, to act as signalling hubs, these integrin nanoclusters should be able to organize other signalling molecules in nanoclusters around it, thereby bringing about downstream signalling. We observe that, other signalling proteins in the focal adhesions including paxillin, FAK, and vinculin form nanoclusters coinciding or in close proximity to integrin β3 nanoclusters [46, 49]. Furthermore, the dynamics of FAK nanoclusters was also observed to be similar to integrin β3, where distinct long-lived nanocluster were observed, suggesting it being organized by integrin β3 nanoclusters and in turn these stable nanoclusters bring about downstream signalling such as force dependent activation of paxillin. Vinculin and paxillin, on the other hand, demonstrated single mixed populations which might be due to their function wherein paxillin, upon activation by force dependent phosphorylation is translocated to the nucleus, or the presence of a free cytoplasmic population, which might allow for transient interactions and faster exchange [18, 59].

Third, to understand if these nanoclusters are signalling hubs, clear signalling events should occur within the nanoclusters. Phosphorylation of proteins including FAK and paxillin is one such event [54–58]. We observe that both phosphorylated FAK and phosphorylated Paxillin were present in high concentration within these nanoclusters with a similar spatial organization as integrin nanoclusters with an intercluster distance of ∼600 nm [52]. An important hallmark of robust signalling is the ability to turn it on and off as needed [10, 84]. When the signal is off (using FAK mutants that are non-phosphorylatable), stable FAK nanoclusters were lost, further suggesting that activated signalling occurs in long-lived immobile nanoclusters that can act as signalling hubs. This loss was also observed when we acutely inhibited FAK (using a small molecule inhibitor PF-562271). If phosphyorylation of a protein acts as an ‘on’ switch for signalling to occur, dephosphorylation would act as an ‘off’ switch [9, 10, 55, 58, 63, 85]. In line with this, we observed that when the phosphatase PTPN12, a force dependent phosphatase, is inhibited, the proportion of FAK nanoclusters that were long lived/immobile increased dramatically. Previous studies using paxillin phosphomimetic mutations also show an increased adhesion size and adhesion turnover [86, 87], while FAK non-phosphorylatable mutants inhibited adhesion formation and increased lifetime [66, 88]. This indicates that immobile fraction organized as nanoclusters is the signalling centre or the signalling hub.

Fourth, this signalling has to be specific to an external signal. Both FAK and PTPN12 are force responsive kinase and phosphatase wherein high mechanical forces activates FAK and inactivates PTPN12 [20–23]. We observed that when force was altered by inhibiting myosin activity, the stability and proportion of immobile clusters of FAK nanoclusters decreased. This clearly shows that stable nanoclusters are signalling sites in response to an external stimulus, in this case, force. If we additionally introduce absence of phosphatase activity (PTPN12 inhibition) to this system, FAK nanoclusters continue to remain stable even in absence of force (treatment with Y-compound which inhibits force transduction), demonstrating that a force responsive ‘off’ signal is required to stop signalling. This further emphasizes that nanoclusters of FAK are signalling hubs formed in response to mechanosignalling. In additional supportive evidence, we observe the following-the geometric arrangement of ligands plays a critical role in focal adhesion formation [30, 41, 89, 90]. We have shown robust adhesion formation (large adhesions) and mechanotransduction (YAP localization to the nucleus) proceeds when ligand is presented on nanodiscs of 100nm diameter (similar to the size of nanoclusters) that are separated by ∼600 nm, whereas robust adhesions don’t form when these nanodiscs are spaced 300 (half the distance) or 1000 (almost twice the distance) nm apart [41]. In this varying geometric arrangement, we observe that stable, long lived, immobile FAK nanoclusters are observed only on nanodiscs spaced 600 nm apart whereas short lived, mobile clusters increases when they are 300 or 1000nm apart (where robust adhesions are not observed), clearly demonstrating that FAK nanoclusters are stable, long lived, signalling hubs formed downstream of external mechanosignalling (both force and geometric arrangement) and they become mobile, short lived non signalling clusters when a robust mechanosignal is absent.

Taken together these results demonstrate that signalling proceeds through the formation of stable, long-lived, immobile nanoclusters, which disassemble when signalling is stopped. Similarly, immobilization of receptors required for signalling has also been shown for other transmembrane receptors such as EGFR [91], B-cell receptors [92, 93], T cell receptor [94], macrophage Toll-like receptor 2 [95]. In addition, it is not single molecule immobilization but a multi-molecular nanocluster immobilization that forms in response to external signalling and is required to organize downstream signalling.

These results build further on the existing archipelago model of FA, in which single integrins were shown to undergo temporary immobilization, now we demonstrate that this happens in nanocluster islands (and get activated), and freely-diffuse outside the islands [31, 46]. It further strengthens the model as we now show that the immobile islands are long-lived in response to an active signalling and can serve as signalling hubs to organize and activate other signalling molecules in signalling hubs. These immobile nanoclusters can then function to respond to local forces, sensing ECM properties, and act as signalling hubs for focal adhesion-related signalling [70, 96–102] that in turn control many aspects of cell behaviour including polarity and migration [103, 104].

## Materials and Methods

### Cell culture

Mouse Embryonic Fibroblasts (MEFs) [1, 30, 105], FAK^−/−^ [106], Vinculin^−/−^ [18], CHO-B2 (expressing a very low level of β3 integrin) [107] cells were used for experiments. Cells were cultured in high glucose Dulbecco’s modified Eagle’s medium (DMEM) with 10% fetal bovine serum, 1% sodium pyruvate, and 1% penicillin-streptomycin in a humidified 37°C and 5% CO2 incubator. Plasmids were transfected by electroporation (Neon transfection system, Life Technologies) according to manufacturer’s protocols. Experiments were carried out 16-24 h post-transfection by spreading cells on fibronectin-coated dishes for the indicated durations. Thirty minutes before the experiments, cells were resuspended using TrypLE express and allowed to recover in Ringer’s solution [108].

### Expression Vectors

Integrin β3-GFP was described before [109]. Integrin β3-PAGFP and Paxillin-mApple were described before [30, 71]. Paxillin-PATagRFP and FAK-PATagRFP were cloned from Paxillin-mApple or FAK-mApple and PATagRFP from PATagRFP-DAAM1 described before [110]. FAK Y397F-PAGFP was created using point mutation in FAK using the FAK-mApple plasmid and cloned into a PAGFP-N1 expression vector. FAK-mApple Vinculin-mApple, Paxillin-mEos2, FAK-tdEos, Vinculin-mEos2, Paxillin mTagBFP2, Paxillin mIFP, PAGFP-N1, and Vinculin PAGFP were gifts from Michael W. Davidson (The Florida State University).

### Antibodies and Inhibitors used

Phosphorylated FAK (Y397) antibody was purchased from Abcam (Cat No. ab81298; 1:200). Phosphorylated paxillin (Y113) antibody was purchased from Abcam (Cat No. ab216652; 1:200). Alexa 647-labeled secondary rabbit antibody was purchased from Invitrogen (Cat No. A-21245; 1:200). MEFs were treated with either 10 µM FAKin (PF-562271, Sigma-Aldrich, Cat. No. PZ0387), 2.5 µM PTPin (PTP LYP inhibitor, Calbiochem, Cat. No. 540217), 10 µM Y-27632 (Sigma-Aldrich, Cat. No. Y0503), or combinations of these drugs during spreading, as indicated.

### Immunofluorescence

Cells were fixed with freshly prepared 4% formaldehyde (Electron Microscopy Sciences) in PBS for 15 min, permeabilized with 0.1% TritonLX-100 in PBS for 20 min at 37 °C, followed by blocking with 1% BSA (Sigma-Aldrich) for 1 h at room temperature. Fixed cells were first incubated with primary antibody overnight at 4°C, followed by secondary antibody incubation for 2 h at room temperature. Between each step, cells were washed thrice with PBS for 10 minutes each on a rocker.

### Microscopy

For single-molecule localization super-resolution imaging, a TIRF microscope (Olympus IX81, Zero Drift Focus Compensator, Dual camera Hamamatsu ORCA-Fusion BT, Objective 100x) was used. PALM imaging of mEos2 and tdEos constructs was performed in PBS. Low intensity illumination at 405 nm was used to activate the fluorophores and 488 or 561 nm laser was used to image (and bleach) the activated fluorophores. Imaging was performed at 100 ms exposure for 7,000-10,000 frames.

SR-confocal imaging was performed using a spinning-disc confocal microscope based on a CSU-W1 Spinning Disk (Yokogawa) with a single, 70-μm pinhole disk and quad dichroic mirror along with an electron-multiplying charge-coupled device (EMCCD) (Princeton Instruments, ProEM HS 1024BX3 megapixels with 30-MHz cascade and eXcelon3 coating) using a Plan Apo 100× NA 1.45 objective lens. The LiveSR module (Roper Scientific France) was used to provide a structured illumination re-scanner that enhance spatial resolution ∼2× by optical deconvolution.

### NanoKymo-FLAP/ NanoKymo-FRAP

For nanokymo-FLAP, peripheral mature FAs were selected and thin strips of adhesions containing only a few pixels were activated using 405 nm laser, and imaged using the respective emission wavelength (488 nm for PA-GFP, and 561 nm for mEos2, PATagRFP or tdEos) at an interval of 3 s for 5 mins. The cells were placed in the microscope chamber at 37°C for 5 mins prior to imaging, to ensure they were in equilibrium and reduce drift. For nanoKymo-FRAP, photobleaching was performed instead of photoactivation.

For analysis, kymographs of the activated thin strips were created using the KymographBuilder plugin in Fiji. Line profiles for high intensity regions in the kymographs were plotted in Graphpad Prism and fitted using ‘One phase decay’ for nanoKymo-FLAP, and ‘Exponential plateau’ for nanoKymo-FRAP. Plateau values (Y_m_) from the nanoKymo-FLAP fits were used as immobile fraction, while plateau values (Y_m_) from the nanoKymo-FRAP fits were used as recovered fraction. Half-times were calculated using the rate constant (K) from the fits using the formula =ln(2)/K.

### Nanopattern functionalization

Nanopatterned Ti on glass substrates were custom-ordered from ThunderNIL srl (Italy). Ti nanopatterns were functionalized by cyclo-RGD (Peptides International, cat. no. PCI-3895-PI), while glass surface was passivated by supported lipid bilayer following a previously published protocol [71]. Briefly, the Ti nanopatterned coverslips were first cleaned by sonicating in 50% isopropanol. Ti was then converted to TiO_2_ by oxygen plasma (ProCleaner Plus, SKU-1062) and incubated with 1-oleoyl-2-[6-biotinyl(aminohexanoyl)]-sn-glycero-3-phosphoinositol-3,5-bisphosphate-ammonium salt (biotin PI(3,5)P_2_; 200 μg/ml) for 20 min (Avanti Polar Lipids, cat. no. 860565). The surrounding glass surface was passivated using 1,2-dioleoyl-sn-glycero-3-phosphocholine (DOPC) Small Unilamellar Vesicles (SUVs) and blocked with 1% casein for 45 min. Finally, the nanopatterns were functionalized by Dylight 650-Neutravidin (Thermofisher Scientific; cat. no. 84607; 10Lmg/mL for 20 min), followed by biotin cyclo-RGD (10Lmg/ml for 30 min).

### Quantitative Image Processing

For nearest neighbour distance calculations, intensity maxima were found using “Find Maxima…” command on Fiji, and the coordinates were analysed using the dsearchn function in MATLAB. An ideal method to measure this would be Fourier Transformation such as FFT functions. However, because focal adhesions are small and discontinuously organized on the plasma membrane, measuring the intercluster distance within the focal adhesions is not possible using FFT as FFT requires large continuous patterns. To circumvent this problem, we use the nearest point search function to handle the edge effect which arises from clusters at the very edge of focal adhesions. This algorithm measures the distance to 1 nearest neighbour and hence clusters on the edge will have the nearest cluster within the focal adhesion and it circumvents the requirement of large patterns for analysis of nearest neighbour. Gaussian fitting of histograms was performed in Origin using ‘Nonlinear Curve Fit-Gauss’ command for single peaks, and ‘Multiple Peak Fit’ for two peaks.

Actin images were used for binary mask generation for cell area measurement and phosphor-FAK images were used to generate binary mask to measure focal adhesion area. For total adhesion area per cell area, all adhesion areas of sizes >0.1 μm^2^ were added and divided by the respective cell area.

Single-molecule localization was performed by Picasso: Localize software using a Gaussian fit mode [111]. The fitted spots were then reconstructed in Picasso, the maxima were found using Fiji, and further analysis was performed using custom MATLAB scripts. FWHM was calculated by analysing line profiles of individual clusters.

### Statistical analyses and reproducibility

All experiments were repeated between two and three times where we measured >10 cells in each repeat. Graphs were plotted using Origin 2019b and statistical analysis was performed using GraphPad Prism 8. Box plots represent interquartile distances and median, the whiskers represent the standard deviation, and the hollow circle represents the mean. Individual data points were plotted along with the box plots with P-values written on top of the graph.

Data from individual groups was first analysed for normal and lognormal distribution using the Shapiro-Wilk normality test. If all groups were normally distributed, a two-tailed t-test was used for 2 groups, and ordinary one-way analysis of variance (ANOVA), followed by the Tukey’s post hoc test was used for more than 2 groups. If all groups were lognormal, the data was transformed to a normal distribution by taking the log and then analysed. If all groups were neither normal nor lognormal, Mann-Whitney test was used for 2 groups, and Kruskal-Wallis test, followed by Dunn’s post hoc test was used for more than 2 groups.

## Acknowledgments

We thank M. W. Davidson (The Florida State University, Tallahassee, FL, USA), for DNA constructs. This work was supported by intramural funds from the Mechanobiology Institute. K.J. was supported by Mechanobiology Institute Graduate Scholarship, R.C. is supported by Singapore National Research Foundation’s CRP grant (No. NRF2012NRF-CRP001-084), and M.P.S. received National Institutes of Health (NIH) grant support related to this project (no. RO1-GM113022). K.J. and P.K. acknowledge funding support from Ministry of Education Academic Research Fund Tier2 (MOE2019-T2-2-014).

## Author Contributions

K.J. & R.C. designed experiments with help from M.P.S. R.C. performed the experiments with assistance from K.J. and R.F.M. K.J. performed image analysis with assistance from R.F.M. R.C. cloned several of the mutant constructs used here and helped establish data analysis protocols. K.J. and R.C. made figures and wrote the manuscript with inputs from all authors.

