## Supplementary Information for "Nano-clusters of ligand-activated integrins organize immobile, signalling active, nano-clusters of phosphorylated FAK required for mechanosignaling in focal adhesions"

**Immobile integrin nanoclusters organize FAK signalling through force-dependent phosphorylation in focal adhesions**

* - Corresponding

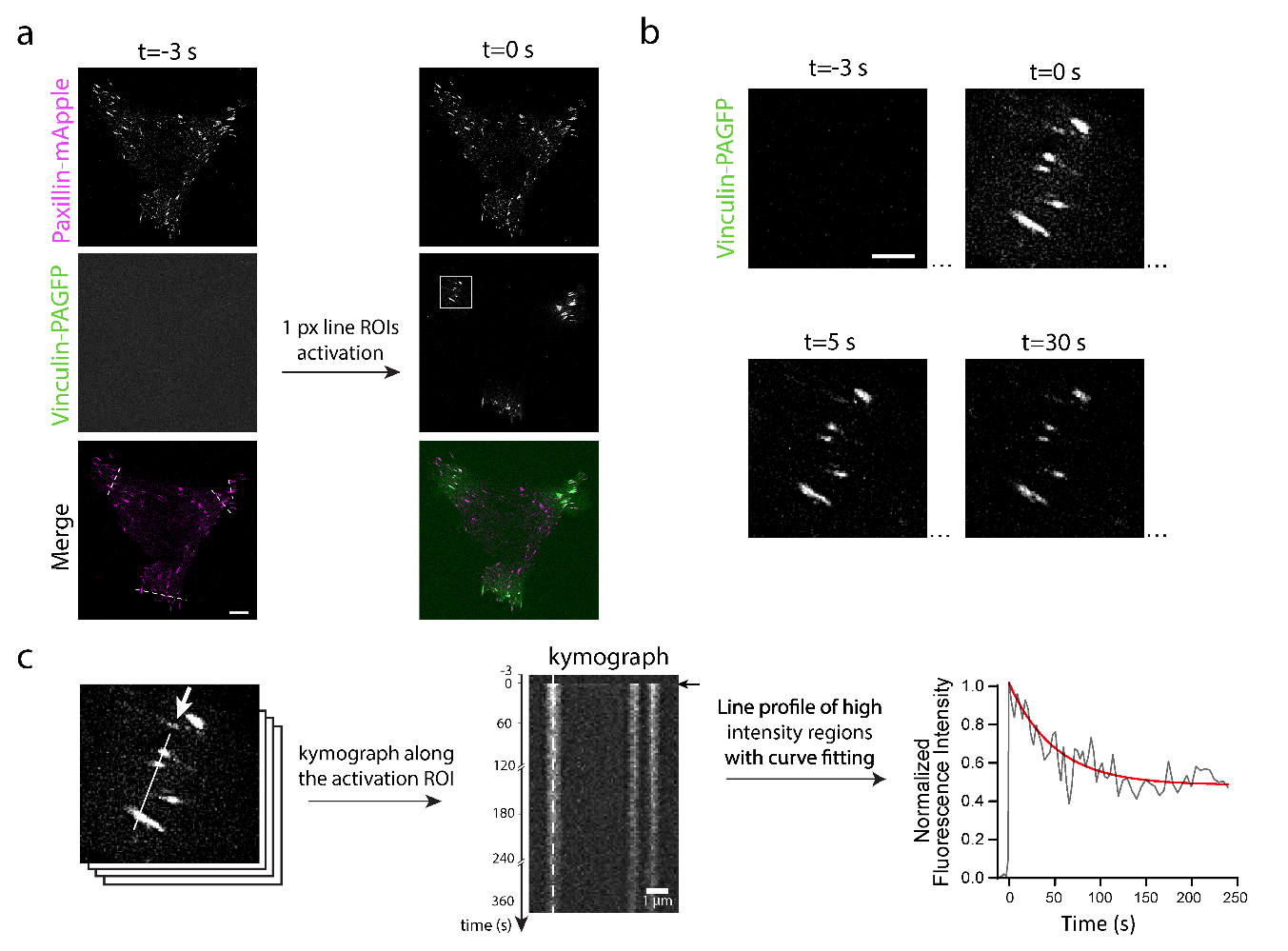

Fig. S1. *Example of the nanoKymo-FLAP technique using MEFs co-transfected with Paxillin-mApple and Vinculin-PAGFP.*

a) Regions of interest (ROIs) of a single-pixel width straight line (along the dashed lines) were activated using PAGFP across multiple focal adhesions and imaged over time. In this example, Paxillin-mApple was used to decide the ROI. Scale bar, 5 µm. b) zoom of the bleached region shown in the white box in a. Scale bar, 2 µm. c) Kymograph along the line ROI was generated and line profiles were created for high-density regions from the kymographs. Scale bar (kymograph), 1 µm. These line profiles were then fitted, and half-time and immobile populations were calculated for the line profile.

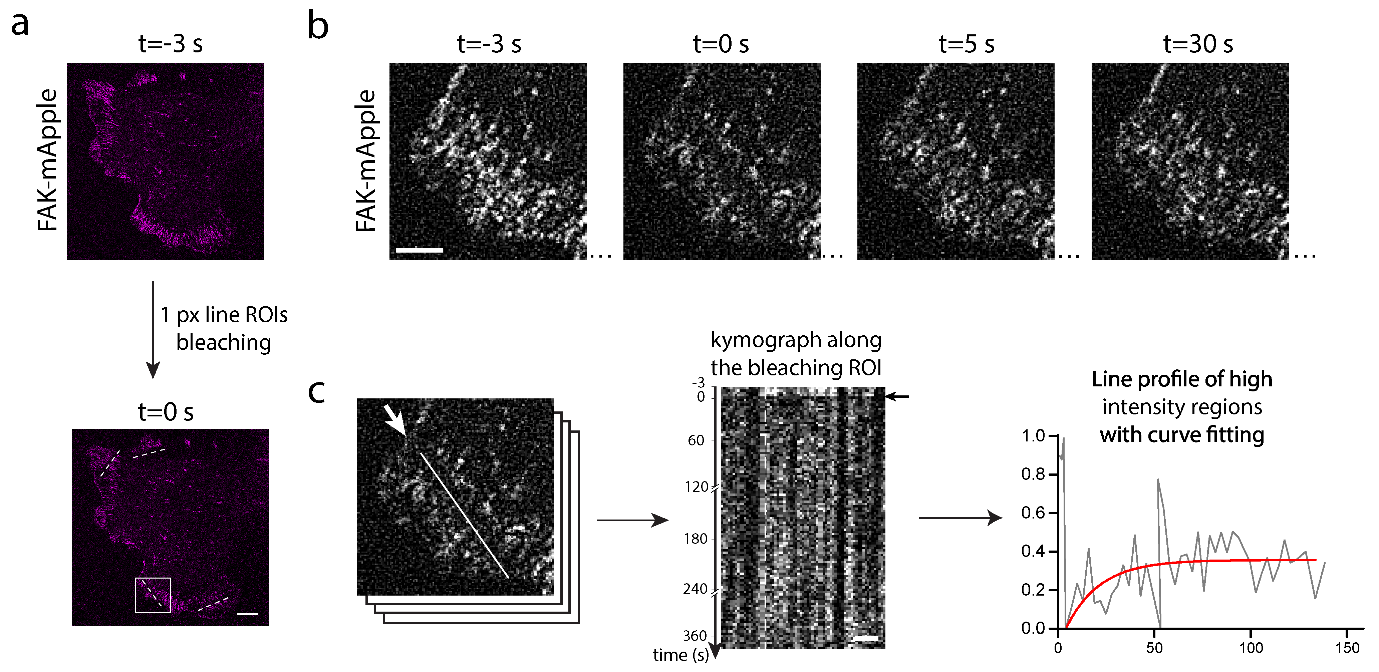

Fig. S2. *Example of the nanokymo-FRAP technique using MEFs transfected with FAK-mApple.*

a) Region of interests (ROI) of a single-pixel width straight-line (white dashed lines) were bleached across multiple focal adhesions and imaged over time. Scale bar, 5 µm. b) zoom of the bleached region shown in white box in a. Scale bar, 2 µm. c) Kymograph along the line ROIs was generated and line profiles were created for high density regions from the kymograph. Scale bar (kymograph), 1 µm. These lines profiles were then fitted, and half-time and recovered population were calculated for the line profile.

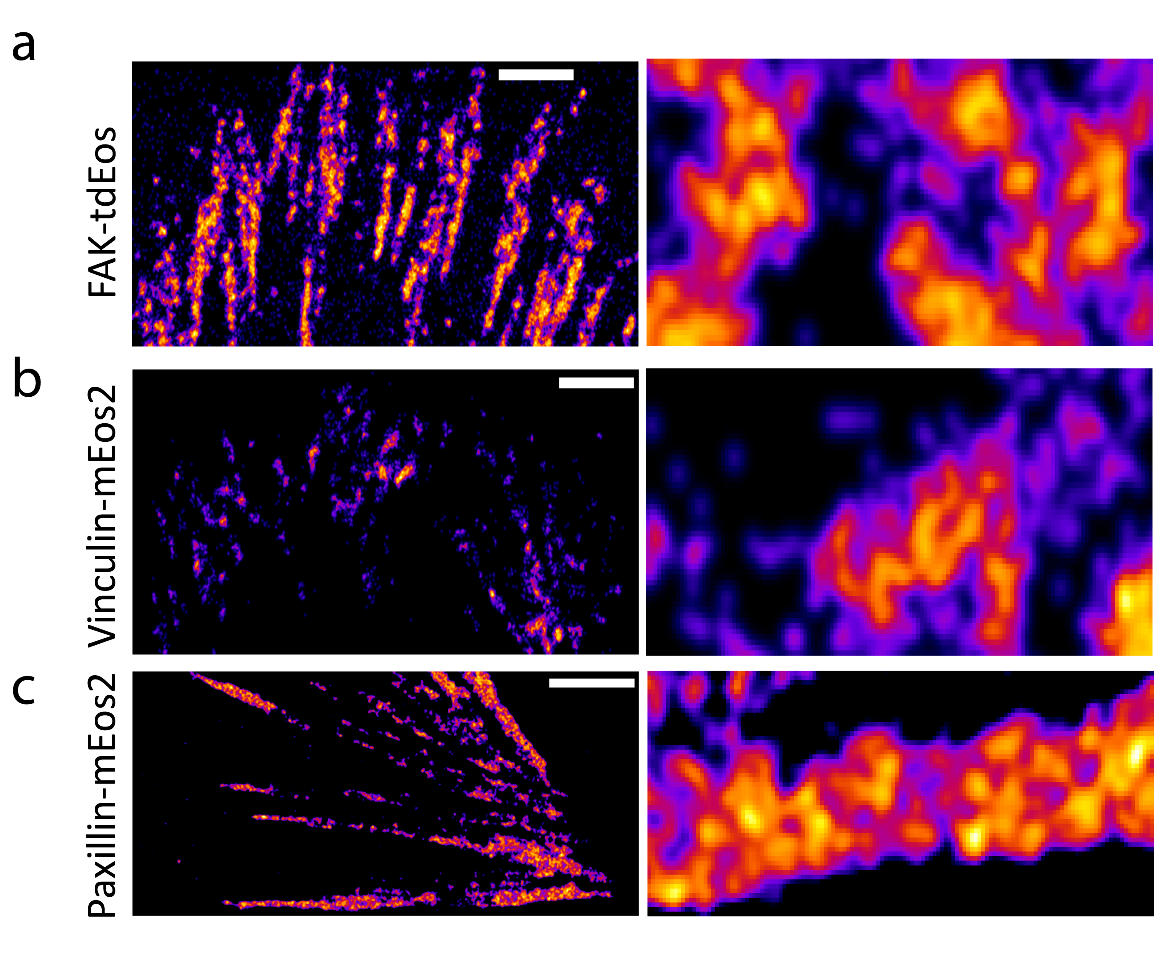

Fig. S3. *PALM imaging of focal adhesion proteins shows nanoclustering.*

Rendered PALM image of (a) FAK, (b) Vinculin, and (c) Paxillin along with the zoom on the right. Scale bar (left) 1 µm, scale bar (right) 500 nm.

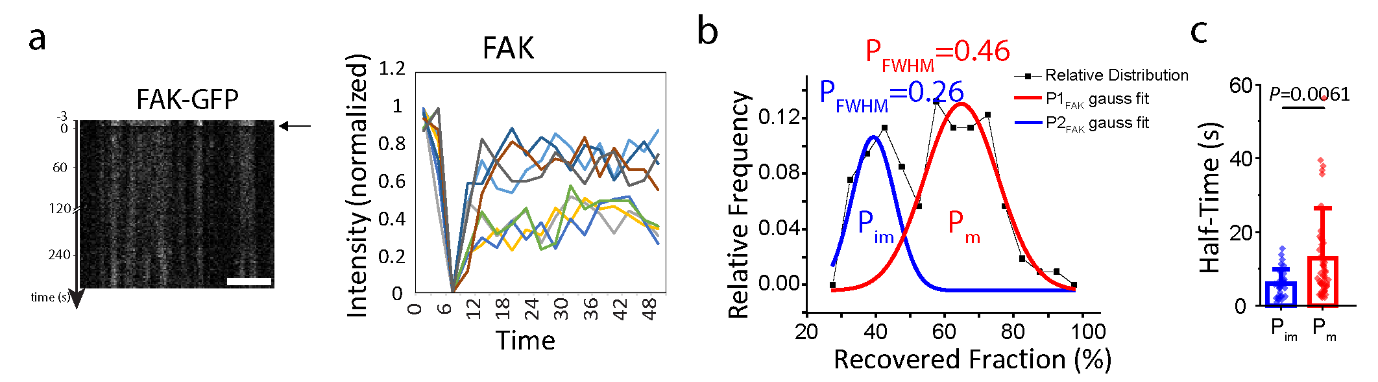

Fig. S4. *NanoKymo-FRAP analysis of FAK shows two distinct populations.*

(a) Representative normalized intensity vs time graphs for nanoKymo-FRAP of MEFs overexpressing FAK-mApple. (b) Relative frequency of recovered populations calculated from fitting intensity vs time plots, along with Gaussian fits of the 2 populations. N=106 from 12 cells. (c) Box plot of half-time fluorescence recovery of the 2 populations. N=32 for P_im_, N=54 for P_m_. Box plots with data overlay display the upper and lower quartiles and a median, the circle represents the mean, and the whiskers denote the standard deviation values, along with the individual points and P values.

Movie S1. Time-Lapse images of MEF overexpressing integrin β3-GFP. Related to Figure 1. Images acquired with super-resolution confocal microscopy. Scale bar, 10 µm.

Movie S2. Time-Lapse images of MEF overexpressing FAK-mApple (left) and integrin β3-PAGFP (right). PAGFP activation was performed at 12 sec. Related to Figure 1.  Images acquired with super-resolution confocal microscopy. Scale bar, 10 µm.

Movie S3. Time-Lapse images of MEF overexpressing Paxillin-mIFP (left) and FAK-PATagRFP (right). PATagRFP activation was performed at 12 sec. Related to Figure 4.  Images acquired with super-resolution confocal microscopy. Scale bar, 10 µm.

Movie S4. Time-Lapse images of MEF overexpressing FAK-GFP. GFP bleaching was performed at 12 sec. Related to Figure S4.  Images acquired with super-resolution confocal microscopy. Scale bar, 10 µm.

Movie S5. Time-Lapse images of MEF overexpressing Paxillin-mApple (left) and Vinculin-PAGFP (right). PAGFP activation was performed at 12 sec. Related to Figure 4.  Images acquired with super-resolution confocal microscopy. Scale bar, 10 µm.

Movie S6. Time-Lapse images of MEF overexpressing FAK-GFP (left) and Paxillin-PATagRFP (right). PATagRFP activation was performed at 12 sec. Related to Figure 4.  Images acquired with super-resolution confocal microscopy. Scale bar, 10 µm.

Movie S7. Time-Lapse images of MEF overexpressing FAK-GFP treated with FAK inhibitor. GFP bleaching was performed at 12 sec. Related to Figure 5.  Images acquired with super-resolution confocal microscopy. Scale bar, 10 µm.

Movie S8. Time-Lapse images of MEF overexpressing Paxillin-mApple (left) and FAK-Y397F-PAGFP (right). PAGFP activation was performed at 12 sec. Related to Figure 5.  Images acquired with super-resolution confocal microscopy. Scale bar, 10 µm.

Movie S9. Time-Lapse images of MEF overexpressing Paxillin-mApple (left) and FAK-PATagRFP (right) treated with PTPN12 inhibitor. PATagRFP activation was performed at 12 sec. Related to Figure 5.  Images acquired with super-resolution confocal microscopy. Scale bar, 10 µm.

Movie S10. Time-Lapse images of MEF overexpressing Paxillin-mApple (left) and FAK-PATagRFP (right) treated with Y27632 at 42 sec. PAGFP activation was performed at 12 sec. Related to Figure 6.  Images acquired with super-resolution confocal microscopy. Scale bar, 10 µm.

Movie S11. Time-Lapse images of MEF overexpressing FAK-GFP. Cells were pre-treated with PTPN12 for 30 mins, imaged for a few frames before Y27632 addition and imaging. Related to Figure 7.  Images acquired with super-resolution confocal microscopy. Scale bar, 10 µm.

Movie S12. Time-Lapse images of MEF overexpressing FAK-mApple spread on nanodisc substrates with specified intercluster distance. Related to Figure 8.  Images acquired with super-resolution confocal microscopy. Scale bar, 10 µm.
